# Epithelial cell extrusion underlies starvation-induced cell loss in a sea anemone

**DOI:** 10.64898/2025.12.12.693960

**Authors:** Inés Fournon-Berodia, Noah Bruderer, Lionel Christiaen, Patrick R. H. Steinmetz

## Abstract

Epithelia likely predate the last common animal ancestor, yet the evolutionary origin and environmental regulation of epithelial remodelling remain poorly understood. Here, we show that extensive, starvation-induced cell loss in the sea anemone *Nematostella vectensis* is associated with epidermal cell extrusion. This process involves formation of a rosette-like arrangement in which an apoptotic, extruding cell is surrounded by a phospho-ERK1/2-positive ring, accompanied by basal translocation of adherens junction components. Combining chemical perturbations with computational quantification of extrusion and cell density, we show that apoptosis is necessary but not sufficient for rosette formation, and that ERK1/2 signalling limits epidermal extrusion density. We further find that extrusion activity increases during starvation, and that extruded cells may be recycled by efferocytosis. Together, our findings indicate that epithelial cell extrusion has physiological roles in sea anemones and that its key hallmarks are evolutionarily ancient, predating the last common ancestor of sea anemones, flies and vertebrates.

## Introduction

Epithelia are a key evolutionary novelty of animals found throughout metazoans, including sponges where their existence was long debated (Leys et al. 2009; Fahey and Degnan 2010). They form a protective barrier and allow the development of structurally distinct tissues and organs (Tyler 2003; Miller et al. 2013). Key molecular components of epithelia, such as the Catenin/Cadherin complex at adherens junctions (AJ) (Fig. 1C, D) or the Integrin-based cell-matrix junction components, are widely conserved across nearly all major metazoan groups (Fig. 1C) (Theocharis et al. 2016; Özbek et al. 2010). Together, these findings led to the commonly accepted hypothesis that epithelia predate the last common metazoan ancestor (Ros-Rocher et al. 2021; Miller et al. 2013; Cereijido et al. 2004). Epithelia undergo a constant cell turnover that is essential for their renewal and integrity, and for responding to environmental insults, such as wounding. An imbalance in this turnover can have dramatic effects leading to over-proliferation or reduced cell loss, which are prime causes of many cancer types (Macara et al. 2014; Eisenhoffer and Rosenblatt 2013; Pothapragada et al. 2022). Keeping epithelial homeostasis is therefore important and achieved by tightly balancing cell proliferation and cell loss, which occurs by apoptosis and epithelial cell extrusion.

**Fig. 1.**
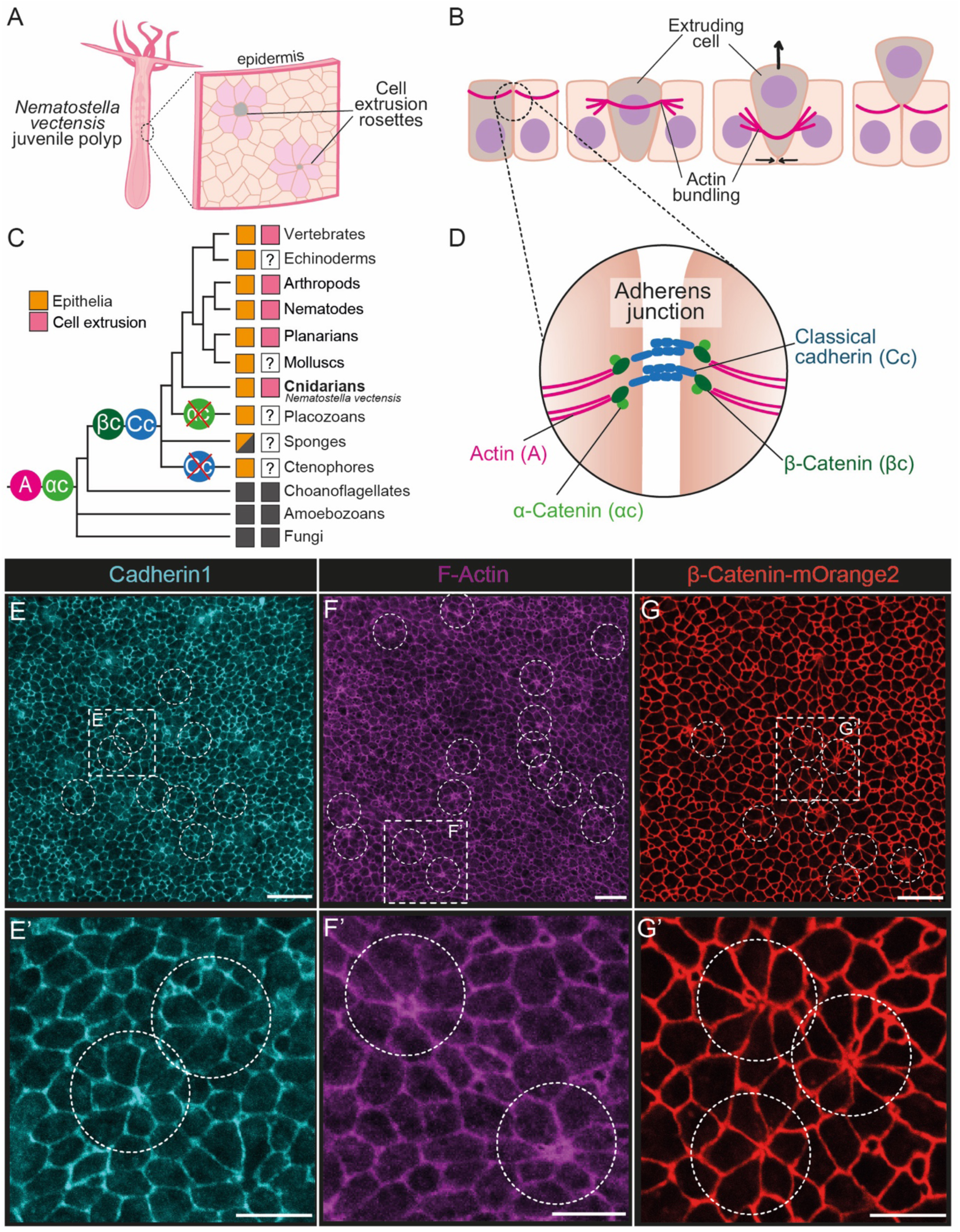
Identification of rosette-like cell arrangements in the epidermis of juvenile *Nematostella vectensis.* (A) Schematic of a juvenile *Nematostella* polyp indicating the body region used for imaging of the epidermis throughout this work. (B) Schematic summary of bilaterian epithelial cell extrusion, focussing on the dynamics of adherens junction (AJ)-associated F-actin bundles (D). From left to right: cell extrusion is driven by the constriction of F-actin (magenta) rings in the extruding cell and adjacent cells, leading to detachment of the extruding cell from the basement membrane. (C) Simplified phylogenetic tree highlighting the phylogenetic occurrence of epithelia, cell extrusion and key AJ components, and the position of Cnidaria (incl. *Nematostella vectensis*) as sister group to Bilateria. Crossed circles indicate secondary losses within taxon. Half grey boxes represent character absence across taxon. (D) Schematic summary of AJ molecules. (E-G’) Projections of confocal imaging z-stacks showing fixed tissue-staining using an antibody detecting *Nematostella* Cadherin-1 (E, E’), Phalloidin to detect F-actin (F, F’), or showing live imaging of β-Catenin-mOrange2 transgenic reporter animals (G, G’). Scale bars: 10 µm (E–G), 20 µm (E’–G’).

Over the last decade, studies in mammals, zebrafish, nematodes and flies have uncovered epithelial cell extrusion as a tightly regulated process to remove single epithelial cells without compromising the structural integrity of the epithelium (Rosenblatt et al. 2001; Gibson and Perrimon 2005; Gu et al. 2011; Eisenhoffer et al. 2012; Marinari et al. 2012; Levayer et al. 2016; Denning et al. 2012). So far, it has been mainly studied in the context of development, during morphogenesis and as response to mechanical tissue stress (Hogan et al. 2009; Leung and Brugge 2012; Grieve and Rabouille 2014; Lubkov and Bar-Sagi 2014; Nanavati et al. 2020; Monier et al. 2015; Ambrosini et al. 2017), but little is known about its regulation by metabolic factors such as nutrient availability. In vertebrates and flies, cells surrounding the extruding cell adopt a characteristic ‘rosette’ formation, accompanied by actomyosin and AJ remodelling in both extruding and rosette cells (Fig. 1A-B). While rosette formation is a hallmark of epithelial cell extrusion, it can also occur during morphogenetic cell rearrangements (Lecuit and Lenne 2007; Guillot and Lecuit 2013). During extrusion, the central cell typically undergoes Caspase-mediated apoptosis.

However, it remains unclear whether apoptosis triggers or instead occurs secondarily, for example as a result of anoikis caused by loss of basal membrane contact (Frisch and Francis 1994; Frisch 2021). Contraction of the ring-like actomyosin system in rosette and central cells lead to apical, basal or lateral extrusion (Fig. 1B) (Gagliardi and Primo 2019; Torres et al. 2017). A hallmark of extrusion in flies and vertebrates is the activity of MAPK/ERK signalling in single or multiple rows of rosette cells, depending on the species and tissues (Aikin et al. 2020; Takeuchi et al. 2020; Gagliardi et al. 2021; Valon et al. 2021). Currently, it is thought that ERK signalling initiates extrusion and protects the surrounding epithelium against pro-apoptotic signals from the extruding cell (Gagliardi et al. 2021; Valon et al. 2021). A major cause of cell extrusion is epithelial crowding, mediated by the mechanosensitive Piezo1 ion channel (Eisenhoffer et al. 2012; Gu et al. 2011). Cell extrusion is also driven by sphingosine-1-phosphate (S1P) that is released from the extruding cell and binds to the S1P receptor in neighbouring rosette cells (Gu et al. 2011; Atieh et al. 2021). How cells are selected for extrusion remains poorly understood, but a recent study showed that cells with low ATP levels are preferentially extruded (Mitchell et al. 2025).

In many bilaterian animals, nutrient availability greatly affects tissue homeostasis, as shown by starvation-induced atrophy of the intestinal epithelium in mice (Chappell et al., 2003; McCue 2012), snakes (Starck and Beese, 2002; Secor 2008), and insects (Park and Takeda 2008; Park et al. 2009; O’Brien et al. 2011; Martin et al. 2018). In extreme cases, starvation leads to whole-body shrinkage that is rare among vertebrates (e.g., fish, amphibians, reptiles) (Bendik and Gluesenkamp 2013; Wikelski and Thom 2000; Field et al. 2007; Huusko et al. 2011; Botius and Botius, 1980), but common among planarians (Baguñà et al., 1990), acoels (De Mulder et al. 2009; Nimeth et al. 2004) and all non-bilaterian taxa, including sea anemones (Chomsky et al. 2004; Sebens 1980; Garschall et al. 2024) (Fig. 1C). So far, however, the environmental control of body and tissue plasticity remains poorly understood on cellular and molecular levels. In *Drosophila*, starvation induces shrinkage of the midgut epithelium by apoptosis- and cell extrusion-mediated cell loss (O’Brien et al. 2011; Martin et al. 2018). Similarly, starvation-induced shrinkage in planarians is mediated by basal cell extrusion in the epidermis, although the underlying cellular and molecular mechanisms remain unclear (Lee et al. 2024; Lindsay-Mosher et al. 2024).

Recently, the sea anemone *Nematostella vectensis* has emerged as a powerful model system for studying the molecular and cellular basis of body plasticity. As in other for cnidarians, *Nematostella* is composed of only two epithelial cell layers: the outer epidermis and inner gastrodermis (Frank and Bleakney 1976; Martindale et al. 2004). Starvation induces substantial cell loss within 2-5 days after feeding, reaching more than 50% after three weeks (Garschall et al. 2024; Pascual-Carreras et al. 2025). Such extensive cell loss necessitates large-scale epithelial rearrangement, making this system uniquely suited to study how environmental cues regulate epithelial remodelling. Here, we show that nutritional fluctuations trigger cell extrusion in *Nematostella*, revealing an ancient epithelial cell elimination mechanism that likely predates the cnidarian–bilaterian split.

## Results

We aimed to identify epithelial cell extrusion in *Nematostella* vectensis (*Nv*) by searching for rosette-like cell arrangements. Cell outlines in the epidermis of the body column were visualised by immunolabelling of three AJ components: NvCadherin1 (Pukhlyakova et al. 2019) (Fig. 1E, E’), F-actin (Fig. 1F, F’), and a β-Catenin-mOrange2 fusion protein in a newly generated transgenic knock-in line (Fig. 1G, G’, S1A). Throughout this study, we focused on the body column epidermis of juvenile polyps, which is particularly accessible to *in vivo* confocal microscopy (Fig. S1B). In all three labelling assays, we observed rosette-like cell arrangements in the *Nematostella* epidermis (Fig. 1E-G’). Because rosettes can also occur during morphogenetic cell rearrangements (Lecuit and Lenne 2007; Guillot and Lecuit 2013), we established *in vivo* confocal time-lapse imaging for the *Nematostella* juvenile epidermis to visualise and characterise cell extrusion events (Fig. 2, S1B). These recordings confirmed progressive constriction and eventual disappearance of the central cell, accompanied by a shift of the intracellular β-Catenin-mOrange2 and F-actin from the apical cortex towards the basal side of the cell (Fig. 2D1-G6, S2 C1-G3). In contrast, the transmembrane AJ component NvCadherin1 remained more apically distributed along the basolateral membrane, absent from the β-Catenin-mOrange2-/F-actin-rich basal ring structure (Fig. S2 J1-N3).

**Fig. 2.**
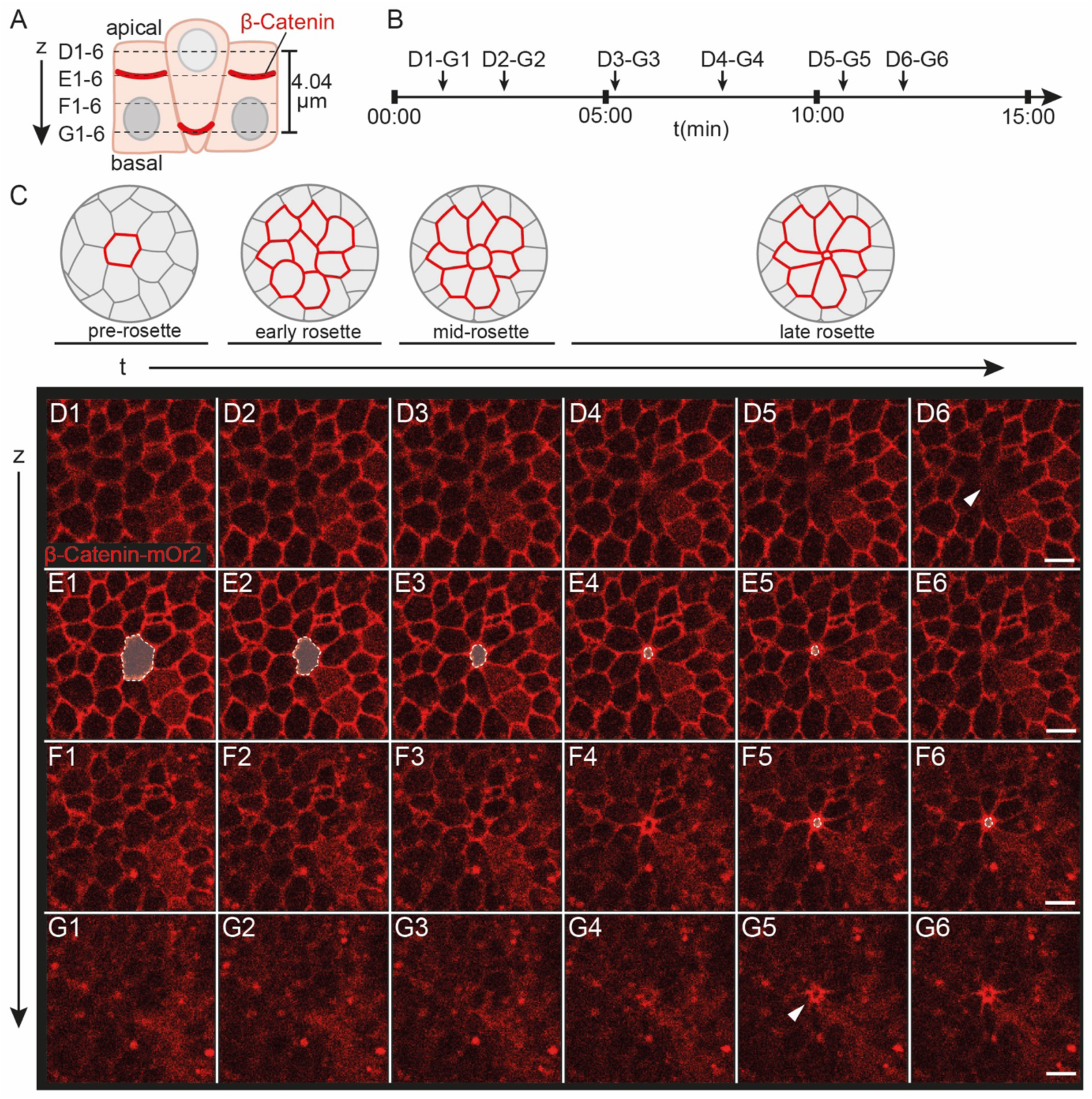
Live imaging of β-catenin-mOrange2 reveals the presence of epidermal cell extrusion in juvenile *Nematostella* polyps. (A, B) Schematics indicating the stack height and relative z-axis positions (A), and the timestamps (B) corresponding to images in (D1-G6). (C) Schematics representing the four stages of rosette formation. (D1-G6) Single plane *in vivo* confocal microscopy images of β-Catenin-mOrange2, depicting the formation of an epithelial rosette from pre-rosette stage (D1-G1) through progressive rosette formation, constriction and extrusion of the central cell (shaded in E1-E5, F5-F6). Note the basal translocation of the central ring (arrowhead). See also Supplementary Video 1. Scale bars: 5 µm.

Based these observations, we propose four stages of epithelial extrusion in the *Nematostella* epidermis: (i) a ‘pre-rosette’ stage (Fig. 2 D1-G1), in which the future extruding cell is indistinguishable from its neighbours; (ii) an ‘early rosette’ stage (Fig. 2 D2-G2, S2 C1-G1), marked by the initiation of rosette formation and central cell constriction; (iii) a ‘mid-rosette’ stage (Fig. 2 D3-G3, S2 C2-G2), during which β-Catenin and F-actin become apparent basally within a clearly defined rosette; and (iv) a ‘late rosette’ stage (Fig. 2 D4-G6, S2 C3-G3), characterised by a basal ring or dot of β-Catenin or F-actin and cumulating in extrusion. *In vivo*, the entire process takes 12 minutes (11:33 ± 2:41mins, n=13 extrusion events) from the early rosette stage initiation to complete extrusion (Fig. S3 B), without clear differences in duration between fed and starved animals (Fig. S3B). This is much shorter than reported for *Drosophila* in which the whole process takes several hours (Martin et al. 2018) and similar to that reported in zebrafish in which it takes approximately 20 mins (Duszyc et al. 2023).

To get insights into the direction of cell extrusion, we studied the location of nuclei within the epithelium. Live imaging of β-Catenin-mOrange2 combined with nuclear staining revealed cases of apical cell extrusion, where the nucleus of an extruding cell appeared apically in the plane of the adherens junction (Fig. S3 A-A’’). The pronounced nuclear DNA stain is consistent with chromatin condensation, one of several hallmarks of apoptosis (Fig. S3 C-C’’) (Voss and Strasser 2020). We also found cases where highly fragmented nuclei, indicative of late apoptosis, locate near the basal side of rosette cells (Fig. S3 D-E’). These may have resulted from basal or lateral extrusion events followed by phagocytosis into rosette cells (see also later results sections). Our observations indicate that epithelial cell extrusion in the *Nematostella* juvenile epidermis occurs primarily apically, but occasional basal or lateral extrusion events remain possible.

Our findings suggest that epidermal cell extrusions contribute to the substantial cell loss reported during starvation-induced body shrinkage (Garschall et al. 2024). We therefore set out to characterise the dynamics of extrusion during feeding and starvation. To quantify epidermal rosettes in a reproducible, high-throughput manner, we developed an image segmentation and rosette prediction pipeline to analyse large numbers of maximum intensity projection (MIP) *in vivo* confocal 2D images of the β-catenin-mOrange2-labelled epidermis in transgenic juvenile polyps (Fig. S4; Methods and Supplementary Methods). Using these images, we segmented individual epidermal cells with a fine-tuned Cellpose model (Pachitariu and Stringer 2022; Stringer and Pachitariu 2025) to automatically predict cell outlines. We then trained a deep learning model (Attention U-Net) to identify candidate rosettes (F1-score = 0.61), which were then verified via a human-in-the-loop workflow. This semi-automated approach accelerated quantification by ∼5-fold compared with manual rosette identification (Fig. S4, Methods and Supplementary Methods). This approach enabled unbiased identification and quantification of cells outlines, cell area and rosettes density across 684 images over both timeseries datasets, during daily feeding (Fig. 3A) and from 1 and 20 days of starvation (Fig. 3B).

**Fig. 3.**
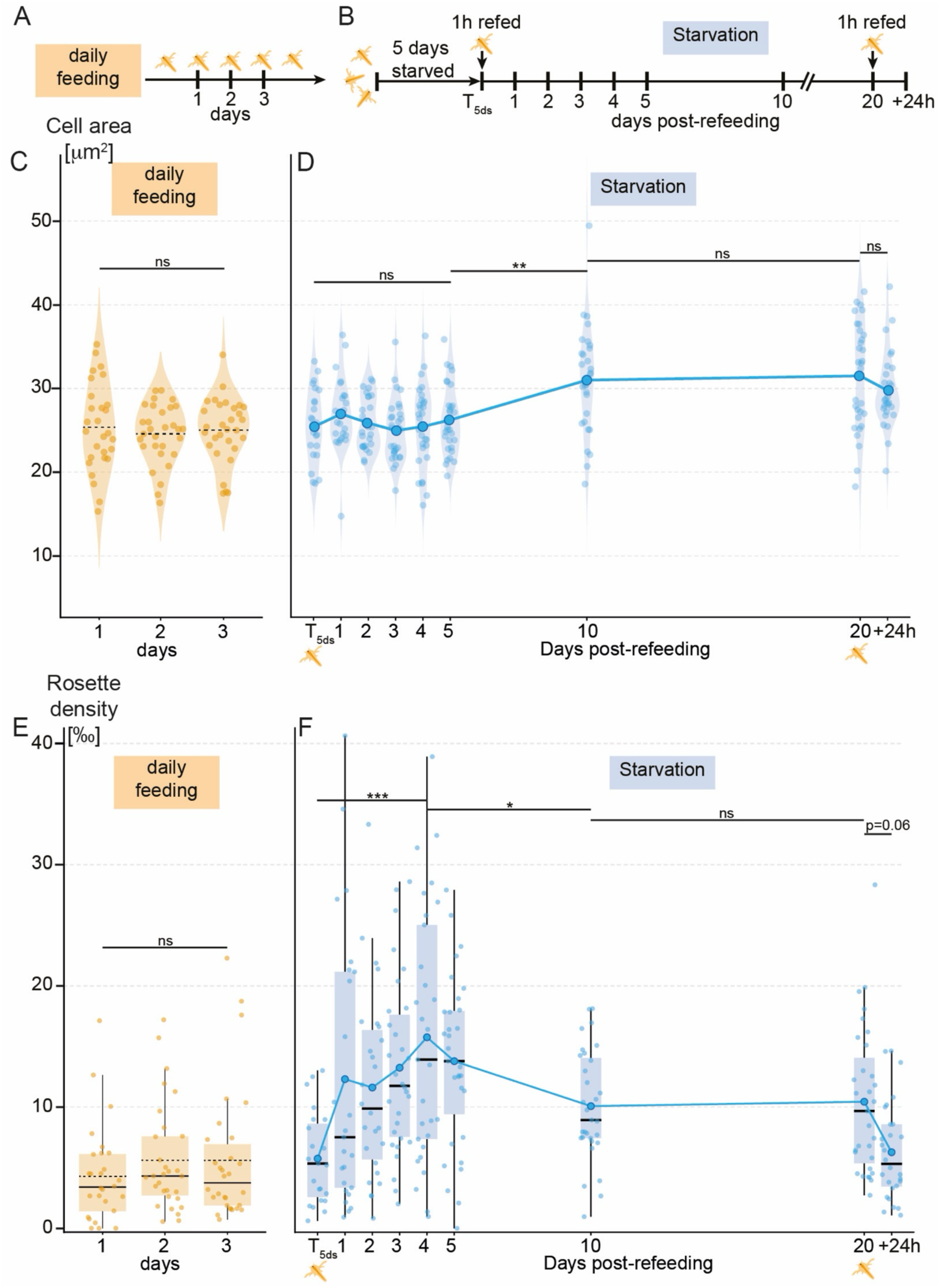
Following rosette density and cell area changes during daily feeding and starvation. (A, B) Sampling strategy for daily feeding (A) and starvation (B) datasets. Labelled vertical stick-marks represent sampled timepoints. (A) Animals were fed daily for 5 hours and sampled over 3 consecutive days. (B) T_5ds_ represents 5 days of starvation after continuous daily feeding. At T_5ds_, animals were refed for 1 hour and then sampled as depicted. At 20 days post-refeeding (dpr), animals were refed for 1 hour and sampled 24 hours later (+24h). (C, D) Violin plots represent cell area distribution of large cells. Means per timepoint are represented as black dashed lines (C) or a central circle (D) that are joined to form a trend line. (C) Mean cell area of large cells is not significantly different across 3 days of daily feeding. (D) Mean cell area of large cells remains stable between T_5ds_ and 5 dpr, significantly increases between 5 and 10 dpr, and remains stable between 10 and 20 dpr. The decrease in mean cell area upon refeeding (+24h) is not significant. (E-F) Box plots depicting rosette density, the number of rosettes identified per 1000 large cells. Means are shown as a black dashed line (E) or as central circles that are joined forming a trend line (F). Median values are shown as black full lines (E, F). Box plots show the first to third interquartile ranges (box), whiskers indicate 1.5x IQR. (E) Rosette density is not significantly different across 3 days of daily feeding. (D) Rosette density increases by ∼2.5-fold between T_5ds_ and 4 dpr, decreases to intermediate levels between 4dpr and 10dpr and stays stables between 10dpr and 20dpr. The decline in rosette density after refeeding at +24h is close to statistical significance. Significance codes for adjusted P-values: * P<0.05, **P<0.01, ***P<0.001; ns = non-significant. Single data points represent mean cell area of large cells (C, D) or mean rosette density (E, F) per animal. Abbreviations: h: hour, ds: days starved.

In *Nematostella*, a feeding pulse triggers a global, transient proliferative burst lasting around 48-72 hours, including in Vasa+/Piwi+ stem-like cell population (Garschall et al. 2024; Pascual-Carreras et al. 2025). Because this rapid, feeding-dependent increase in cell numbers could influence cell density, known to regulate cell extrusion rates in both vertebrates and flies (Kocgozlu et al. 2016; Levayer et al. 2016), we carefully controlled feeding history in all experiments. Following previous work, we either sampled polyps during daily feeding or initiated starvation with a 1-hour feeding pulse after 5 days of prior starvation (T_5ds_), thereby inducing a controlled, relatively synchronised proliferation response before proliferation returns to baseline during starvation (Pascual-Carreras et al. 2025) (Fig. 3A-B).

Using cell segmentation masks, we quantified epidermal cell area as a proxy for tissue density. We found a bimodal distribution of cell area measurements into two populations (Fig. S5 A): (i) ∼25% small cells of ∼6.3 µm^2^ (95% CI = 6.19-6.43)), likely including cnidocytes and sensory neurons; and (ii) ∼75% larger cells measuring ∼27.0 µm^2^ (95% CI = 26.2-27.9), likely consisting predominantly of epidermal cells (Fig. S5 B-C). The population of small cells showed no significant changes in mean cell area after feeding or during starvation, consistently remaining within a narrow range (6.12–6.78 µm²) (Fig. S5 D-G). We therefore focussed our analysis on changes in the population of larger epidermal cells (Fig. 3C-D). Within this population, mean cell areas did not significantly differ between any time points over three days of daily feeding (Fig. 3C), between T_5ds_ and 5 days post-refeeding (dpr) or between 10 and 20 dpr (Fig. 3D). However, we found a significant, ∼20% increase in mean cell area between 5 and 10 dpr (Tukey-adjusted p < 0.001), from ∼26 µm² (95% CI: 24.38–27.62) to ∼30µm² (95% CI: 28.55–32.36) (Fig. 3D). Notably, cell areas at 10 and 20 dpr (∼31 µm², 95% CI 29.36-34.01) are also significantly higher (∼27% adjusted p-value <0.001) compared to daily fed samples (∼25µm², 95% CI 23.31-26.76). Together, this data shows that cell density is similar between daily fed and over 5 days after refeeding at T_5ds_, but that the epidermis becomes significantly less dense during prolonged starvation.

Across all timeseries datasets (Fig. 3A, B), epidermal rosettes typically consisted on average of 6.00±1.82 cells, nearly identical to the number reported in the mammalian gut (∼6 cells) (Simmons et al. 2015). We then quantified epithelial rosette density, corresponding to the ratio of rosette numbers per 1000 large cells in the body column epidermis. In daily fed polyps, we detected ∼5‰ epidermal rosettes (Fig. 3E; 95% CI = 3.42-6.91). Rosette density was higher at T_5ds_ (∼6‰; 95% CI = 1.26-10.58) and continued to increase significantly to ∼15‰ at 4 days post-refeeding (dpr) (95% CI = 10.21-19.87; Tukey-adjusted p < 0.001), constituting a ∼1.5-fold increase (Fig. 3F). Between 4 and 10 dpr, the mean rosette density decreased significantly (Tukey-adjusted p=0.033) by ∼30% to intermediate levels of ∼10‰ (95% CI = 5.20-15.18) (Fig. 4F). After a 1-hour refeeding pulse after 20 days of starvation (20dpr+24h), rosette density tended to decline nearly significantly (Tukey-adjusted p=0.0592) by ∼40% to ∼6‰ (95% CI = 1.28-11.52), within 24 hours, suggesting that refeeding at 20dpr has opposite effects on extrusion dynamics compared to refeeding at T_5ds_ (Fig. 4F). Notably, rosette density at 10 (∼10‰; 95% CI 7.42-12.64) or 20 dpr (∼10; 95% CI 8.1-13.33) are still ∼2-fold higher than compared to daily fed animals (∼5‰, 95% CI 2.59-8.19) (Fig. 3 E-F, Suppl. Table 5). In summary, our results reveal a parallel trend in tissue density and extrusion dynamics. Cell extrusion levels are generally low during continuous feeding, when polyps are dynamically growing. Notably, refeeding after 5 days of starvation lead to a pronounced increase in cell extrusion events at relatively high cell density, followed by a coordinated decline in extrusion and cell density beyond 4 dpr. During prolonged starvation, rosette density stays at intermediate, but still considerably higher, levels than under daily feeding.

**Fig. 4.**
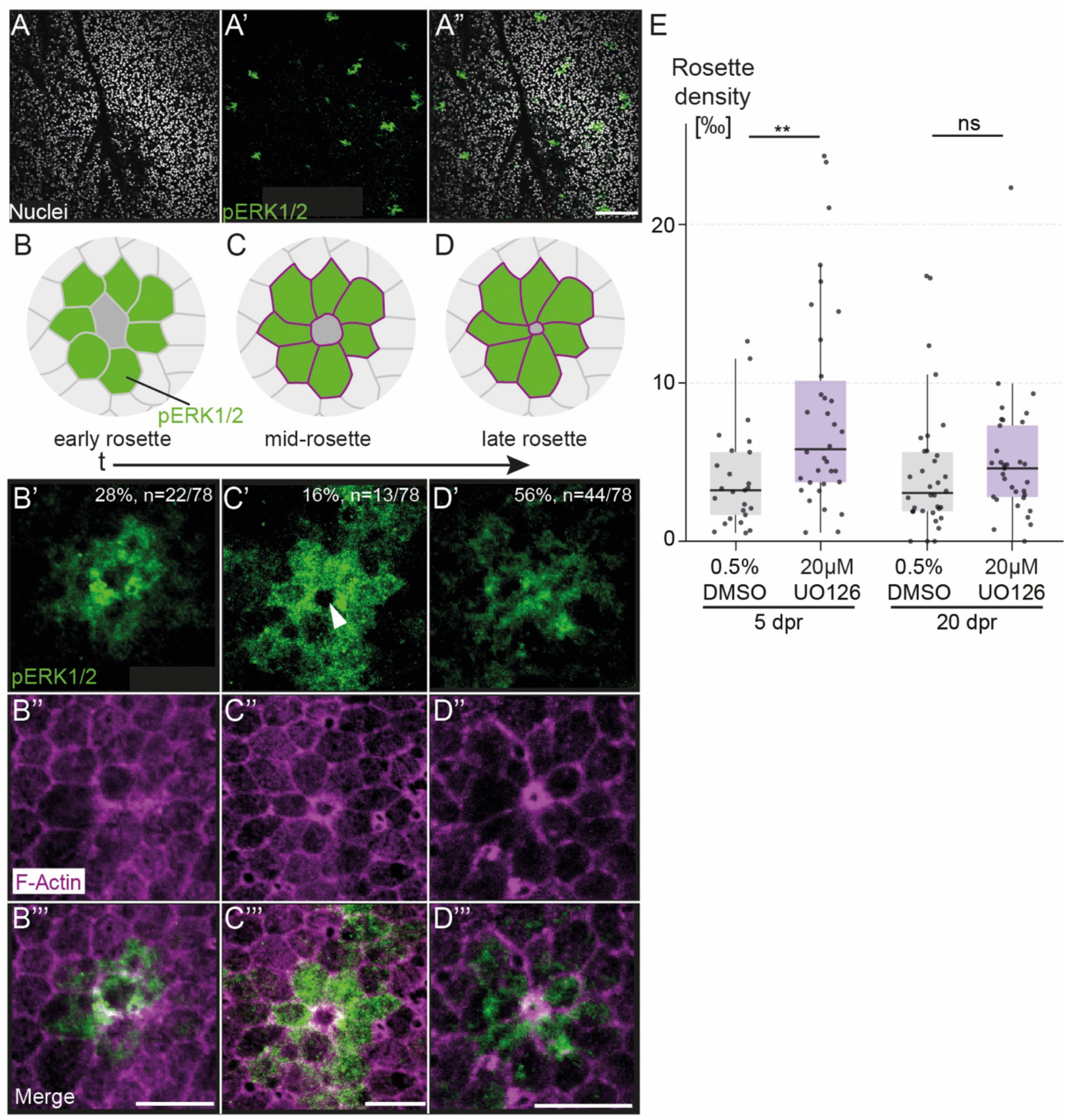
ERK1/2 signalling is activated in rosettes, and its inhibition increases rosette density. (A–A’’) Projections of confocal microscopy images showing scattered cell patches immunolabelled by phospho-ERK1/2 (pERK1/2) in the fixed juvenile *Nematostella* epidermis. (B-D’’’) Schematics (B-D) summarizing the results of Z-projections of confocal microscopy images depicting co-localisation of pERK1/2 label (B’, B’’’, C’,C’’’ D’, D’’’) with F-actin-labelled rosettes (B’’, B’’’, C’’, C’’’, D’’, D’’’). (E) Box plots show mean rosette density (i.e., rosette number per 1000 large cells) in the epidermis of polyps at 5 or 20 days post-refeeding treated for 2 hours in the MEK inhibitor U0126 or in 0.5% DMSO as control. Inhibition of MEK, the kinase directly upstream of ERK1/2, led to an increase in rosette density that was only significant at 5dpr. Box plots show median values (black full lines), the first to third interquartile ranges (box), whiskers indicate 1.5x IQR. Single data points represent mean rosette density per animal. Significance codes for model adjusted P-values: ** = P<0.01, ns = non-significant. Abbreviations: dpr: days post-refeeding. Scale bars: 50 µm (A) and 10 µm (B-D).

To gain mechanistic insights into cell extrusion in *Nematostella* and enable molecular comparisons with bilaterians, we studied the role of ERK/MAPK signalling during extrusion (Fig. 4, S6, S7). In vertebrates and flies, the ERK/MAPK pathway is induced in rosette cells, coordinates the spatiotemporal dynamics of extrusion and protects the surrounding epithelium from pro-apoptotic signals (Gagliardi et al. 2021; Villars and Levayer 2022). We tested whether ERK/MAPK activity in rosettes is evolutionary conserved in *Nematostella* by performing immunolabelling of phosphorylated ERK1/2 (pERK1/2; Fig. 4 A-D’’, S6 A-C’’’) using a commercial antibody previously validated in *Hydra* (Tursch et al. 2022). Using this antibody, we found pERK1/2+ cell clusters broadly co-localising with early to late-stage rosettes throughout the epidermis (Fig. 4 A-D’’’). Manual quantification showed that ∼89% of all rosettes identified by F-actin labelling were pERK1/2+ (n=78/88; Fig. 4B’, C’, D’ and Supp. Table 26). These observations support a strong association between ERK1/2 signalling activity with rosette formation beyond early stages. While the central, extruding cell lacked any detectable pERK1/2 signal (Fig. 4C’, white arrowhead), the outer pERK1/2 signal boundary was often not sharply delimitated, with weaker labelled cells found beyond the rosette cells (f.ex.: Fig. 4C’).

To assess the function of ERK during cell extrusion, we used the small molecule U0126 to inhibit the mitogen-activated extracellular signal-regulated kinase (MEK), which is directly upstream of ERK1/2 and validated previously in *Nematostella* larvae (Favata et al. 1998; Rentzsch et al. 2008). We confirmed that a 2-hour treatment with 20 µM U0126 effectively decreased pERK levels below the detection limit also in the juvenile epidermis (Fig. S7 A-B’). Under these experimental conditions, we found that MEK inhibition at 5dpr, when cell and rosette densities are high, caused a significant increase in rosette density compared to the 0.5% DMSO-treated control (+88% change, adjusted p-value = 0.0027) (Fig. 4E). At 20dpr, when cell and rosette densities have declined, MEK led only to a mild, non-significant increase in mean rosette density (by ∼14%; p = 0.57) (Fig. 4E). In contrast to rosette density, the mean cell area of large epidermal cells was not significantly different between U0126- and DMSO-treated polyps at 5 or 20 dpr (Fig. S7 C). We thus conclude that the U0126-induced increase in rosette density is not indirectly caused by changes in tissue density. Our findings thus indicate that ERK/MAPK signalling regulates cell extrusion density independently of tissue density in the *Nematostella* epidermis. Together, our results highlight ERK/MAPK signalling activity in rosette cells as an evolutionary conserved feature of cell extrusion between *Nematostella*, flies and vertebrates.

To characterise the role of apoptosis during *Nematostella* cell extrusion, we tested morphological and molecular markers that have been associated with different stages of apoptosis in bilaterians (Zmasek and Godzik 2013). We first studied p38/MAPK signalling, which has been linked to apoptosis in *Drosophila*, planarians and mammals (Santabarbara-Ruiz et al. 2015; Arnold et al. 2016; Deschesnes et al. 2001; Yue and López 2020). Using a phospho-specific p38 antibody (p-p38) previously validated in the cnidarian *Hydra* to detect p38 activity (Tursch et al. 2022), we found that p-p38+ cells often exhibited condensed or fragmented nuclei, indicative of late apoptotic states (Fig. S6 A-A’’). Co-staining against pERK1/2 antibody furthermore confirmed that at least some p-p38+ cells were surrounded by pERK1/2+ cells within an extrusion rosette (Fig. S6 B-C’’’). To capture earlier stages of apoptosis, we used an antibody detecting the conserved Caspase cleavage substrate motif (CCS) that become exposed upon Caspase-catalysed proteolysis (Xu et al. 2019; Ma et al. 2019; Stokes et al. 2016; Mazloumi Gavgani et al. 2025). After aligning fixed, F-actin-stained rosettes along the proposed staging system, we found CCS+ central rosette cells throughout all stages starting at pre- or early rosette stage (Fig. 5A-C’’’). At the earliest stage, CCS+ cells were of comparable size to their neighbours, making it difficult to determine if rosette constriction had already been initiated (i.e., early rosette) or not (i.e., pre-rosette) (Fig. 5A-A’’’). At late rosette stages, we observed that some extruded CCS+ cells were retained at the epithelial surface (Fig. 5F-F’’). Manual quantification revealed that ∼81% of F-actin-defined rosettes (n=46/57) contained a central CCS+ cell (Fig. 5’, B’, C’ and Suppl. Table 26), suggesting that extruding cells predominantly undergo apoptosis during the process.

**Fig. 5.**
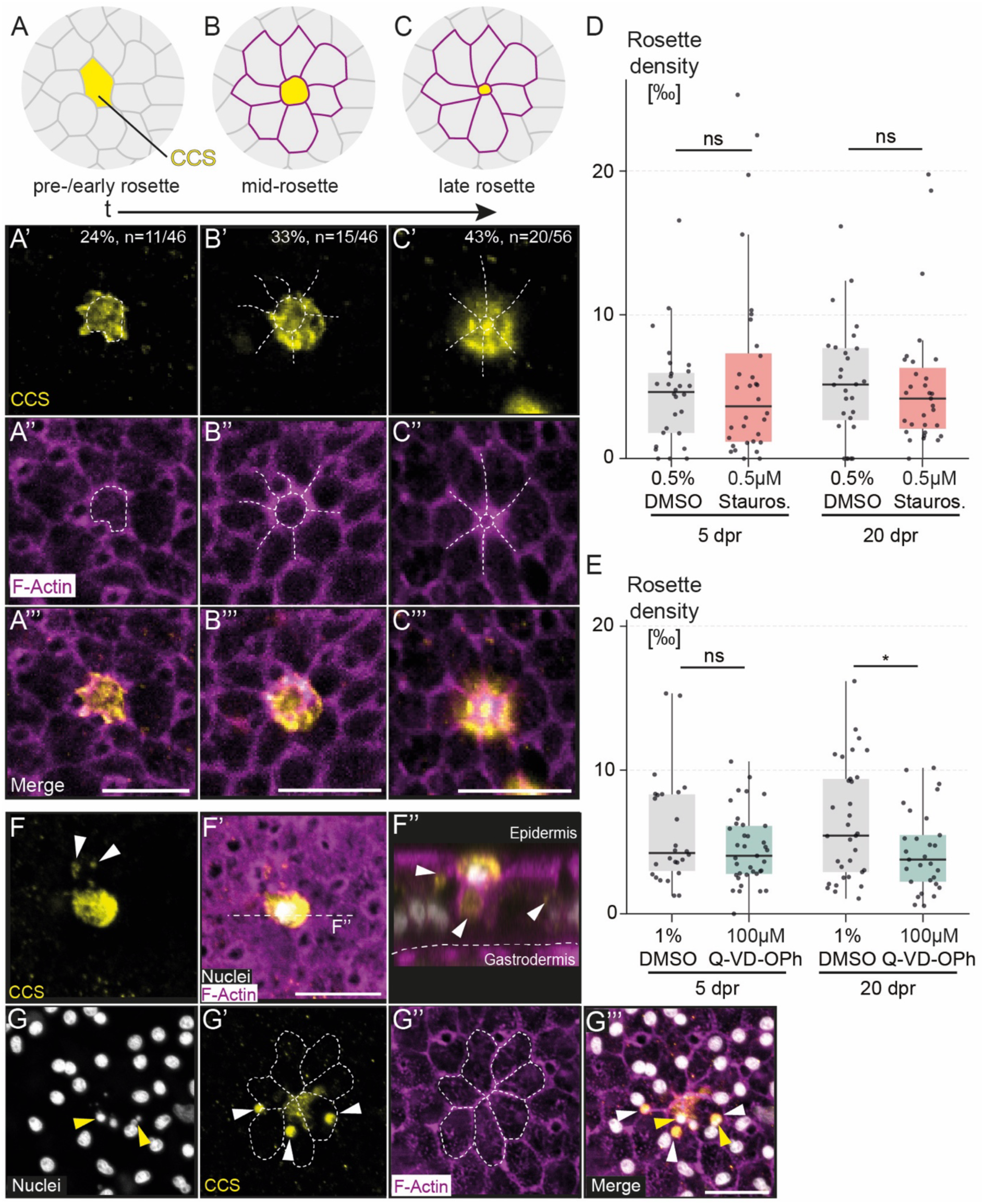
Caspase-mediated apoptosis is a hallmark of the extruding cell. (A-C’’’) Schematics (A-C) summarizing the results of projections of confocal microscopy images depicting localisation of Cleaved Caspase substrate (CCS)-immunolabelled cells (A’, A’’’, B’, B’’’, C’, C’’’) within the central cell of an F-actin-labelled rosette (A’’-A’’’, B’’-B’’’, C’’-C’’’). (D-E) Box plots show mean rosette density (i.e., rosette number per 1000 large cells) in the epidermis of polyps at 5 or 20 days post-refeeding treated for 2 hours in the apoptosis inducer Staurosporine (D), the pan-Caspase inhibitor Q-VD-OPh (E) or in DMSO as control (D, E). (D) Induction of apoptosis by Staurosporine had no significant effect on rosette density at either timepoint. (E) Pan-caspase inhibition caused a decreasing trend in rosette density at both timepoints tested, which was only statistically significant at 20dpr. (F–F’’) Projections (F, F’) and z-axis reconstruction (F’’) of confocal imaging stack of CCS/F-actin co-labelled epithelial cells show apically extruded CCS+ cell, and presence of putative phagocytic vesicles containing CCS+ apoptotic cell debris in adjacent cells (white arrowheads). Dotted line in F’ depicts the plane of reconstruction in F’’. (G–G’’’) Projections of confocal imaging stack of CCS/F-actin/nuclear stain co-labelled epithelial cells showing presence of putative phagocytic vesicles containing CCS+ (white arrowheads), including nuclear fragments (yellow arrowheads), in adjacent cells. Significance codes for model adjusted P-values: * P<0.05, ns = non-significant. Box plots show median values (black full lines), the first to third interquartile ranges (box), whiskers indicate 1.5x IQR. Abbreviations: dpr: days post-refeeding, Stauros.: Staurosporine. Nuclear stain: Hoechst. All scale bars: 10 µm.

These observations prompted us to study the role of apoptosis during cell extrusion using Staurosporine, inducing apoptosis by non-selective inhibition of protein kinases (Pothapragada et al. 2022; Chae et al. 2000; Zhang et al. 2004). We confirmed that 0.1-1µM Staurosporine consistently induces apoptosis in the *Nematostella* body column epidermis in a concentration-dependent manner (Fig. S8 A-D’). Incubating polyps at 0.5µM Staurosporine for 2 hours, which triggers intermediate levels of apoptosis induction (Fig. S8 C, C’), led to no significant changes in rosette density compared to 0.5% DMSO-treated controls at 5dpr or 20dpr (Fig. 5D). In contrast, Staurosporine caused a decrease in mean cell area at both 5dpr (model adjusted p-value = 0.09) or 20dpr (model adjusted p-value = 0.043) that may result from cell loss by apoptosis (Fig. S8 G). Additionally, Staurosporine-mediated apoptosis causes a broad increase in ERK activity across the epidermis, likely as stress response (Sun et al. 2015; Gagliardi et al. 2021; Crozet and Levayer 2023) (Fig. 5E-F’). These findings show that apoptosis-mediated cell loss may occur independently of extrusion, and that the induction of apoptosis is not sufficient to initiate epidermal cell extrusion.

To further explore whether apoptosis is essential for the initiation or progression of cell extrusion in *Nematostella*, we used the pan-Caspase inhibitor Q-VD-OPh to test whether blocking apoptosis affects rosette density (Caserta et al., 2003; Marchiando et al. 2011). Using confocal imaging, we confirmed that incubation in 100µM Q-VD-OPh over 2 hours robustly decreased epidermal CCS immunolabelling compared to 2% DMSO-treated controls (Fig. S8 H-I’). Under the same conditions, we found the inhibition of apoptosis tends to suppress rosette density at both 5dpr (-20%, model-adjusted p-value = 0.17) and 20dpr (-31%, model-adjusted p-value = 0.018) (Fig. 5E). Cell density, however, was not significantly affected between Q-VD-OPh and DMSO controls at any timepoint (Fig. S8 J). Together, our results suggest that inducing apoptosis is not sufficient to trigger epithelial cell extrusion in the *Nematostella* epidermis, but that it appears to be necessary at least in some cases for extrusion formation.

During late rosette stages, we occasionally observed CCS+ intracellular vesicles of 1-2µm located within rosette cells (Fig. 5G-G’’’ white arrowheads). These observations indicate efferocytosis, the engulfment of apoptotic material, by neighbouring rosette cells. It also intriguingly suggests that cell extrusion may contribute to nutrient recycling during starvation. To test this assumption, we studied TOR signalling activity, a nutrient-sensitive regulator of anabolic processes (e.g., protein translation rates) conserved across fungi and animals (Cargnello et al. 2015; Laplante and Sabatini 2012). Active TOR signalling was detected by immunolabelling of phosphorylated ribosomal S6 protein (pRPS6), a widely conserved TOR signalling readout previously validated in *Nematostella* (Ikmi et al. 2020; Voss et al. 2023; Pascual-Carreras et al. 2025). As expected, we found broad epidermal pRPS6 staining in juvenile polyps at 24 hours after feeding (Fig. 6A-A’’). After 20 days of starvation, however, the pRPS6 signal was confined to distinct epidermal clusters overlapping almost specifically with late rosette stages (Fig. 6B-D’’). pRPS6 immunolabel was almost specific to cells of late rosette stages (n=18/19). Some of these CCS+ rosette cells contained nuclear fragments reminiscent of the CCS+ vesicles (Fig. 6D, yellow arrowheads). Together, these results suggest a model in which efferocytosis of extruded apoptotic cells by rosette cells contributes to nutrient recycling under starvation. Alternatively, TOR signalling could play a mechanistic role in rosette formation, for example to regulate the actomyosin cytoskeleton as previously described for TOR complex 2 (Jacinto et al. 2004; Baker et al. 2016). We tested this possibility by studying how AZD8055, a small molecule inhibitor of TOR complexes 1 and 2, affects extrusion density (Chresta et al. 2010; Voss et al. 2023; Pascual-Carreras et al. 2025) (Fig. S9 A-D). We found that incubating juvenile polyps in 1µM AZD8055 for 2 hours strongly decreases epidermal pRPS6 levels (Fig. S9 A-B’) but did not significantly affect rosette or tissue densities compared to 0.5% DMSO controls (Fig. S9 C-D). Altogether, these data indicate that TOR signalling is not necessary for rosette formation and strengthens the assumption that TOR activation in late rosettes results from nutrient recycling.

**Fig. 6.**
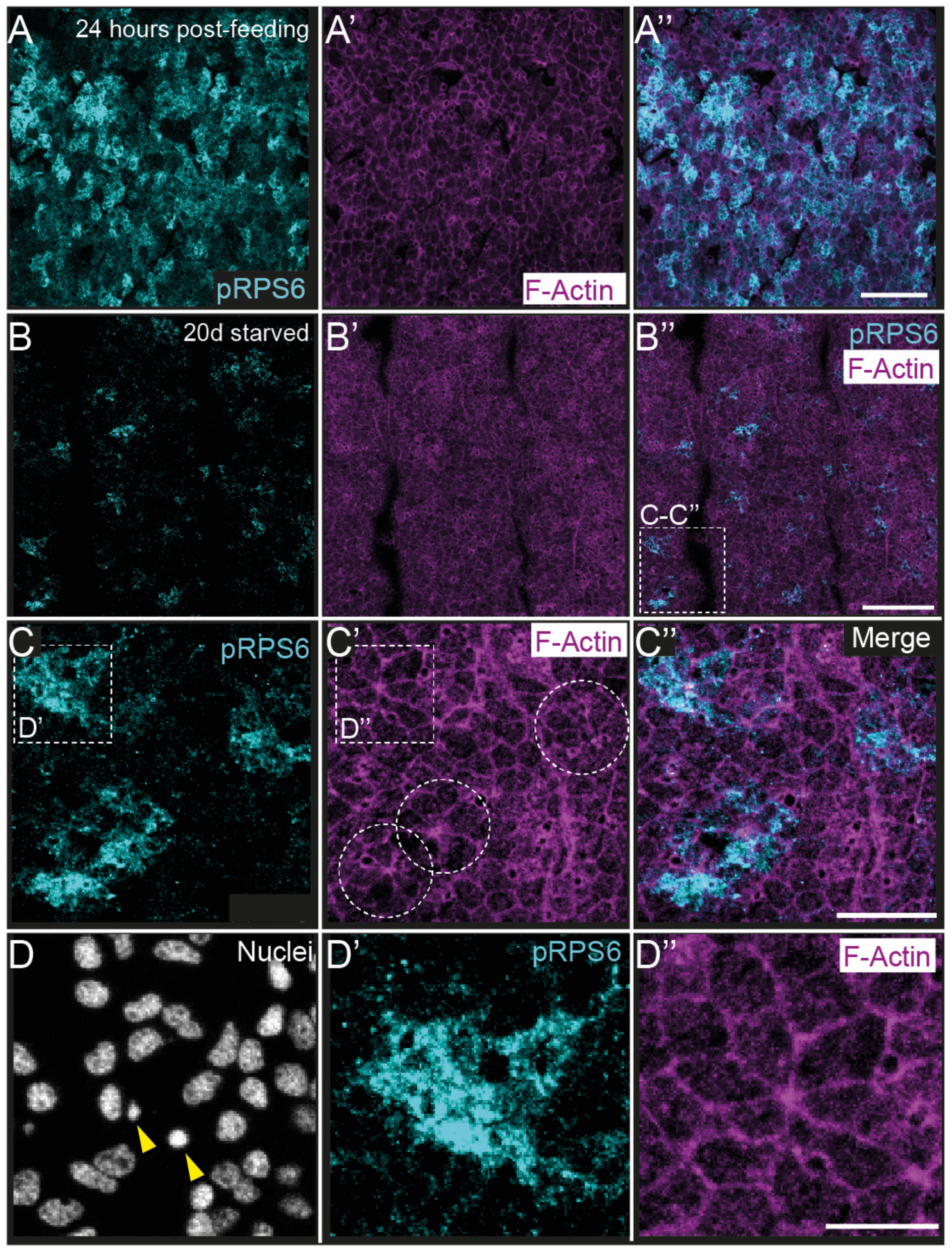
TOR signalling activation in epidermal rosettes of starved polyps. (A-D’’) Projections of confocal imaging stacks of fixed epidermis labelled for phosphorylated Ribosomal Protein S6 (pRPS6), F-actin and nuclei in polyps at 24 hours (A-A’’) or 20 days post-feeding (B-D’’). While pRPS6 signal, a readout for TOR signalling activity, is broadly distributed across the epidermis of juvenile *Nematostella* polyps (A-A’’) at 24 hours post-feeding, it is restricted to late extrusion rosettes after 20 days of starvation (B–D’’). Notably, some pRPS6+ cells contain condensed nuclear fragments (yellow arrowheads), reminiscent of the CCS+ vesicles (Fig. 5G-G’’’). Nuclear stain: Hoechst. Scale bars: 50 µm (B), 20 µm (A, C) and 10 µm (D).

## Discussion

Epithelia are found across all animal taxa, yet the evolutionary origins of their plasticity and their regulation by environmental cues remain poorly understood. In this study, we used the epidermis of the sea anemone *Nematostella vectensis* as a tractable system to investigate the cellular mechanisms underlying starvation-induced cell loss and body shrinkage. Juvenile polyps lose more than half of their cells within three weeks of starvation, whereas refeeding triggers a pronounced proliferative burst from previously quiescent cells (Garschall et al. 2024; Pascual-Carreras et al. 2025). These observations prompted us to investigate how the epithelium accommodates such extensive cell loss without compromising epithelial integrity. In vertebrates and *Drosophila*, epithelial cell extrusion plays a central role in maintaining epithelial homeostasis and integrity (Krueger et al. 2025; Martin et al. 2018). Using live imaging of a transgenic knock-in reporter line expressing the β-Catenin-mOrange2 fusion protein to visualise AJs, we discovered and characterised epithelial cell extrusion in the *Nematostella* epidermis.

Together with F-actin labelling and immunostainings against Cadherin-1 and pERK1/2, we found that extrusion in *Nematostella* involves the formation of pERK1/2+ rosette cells, the basal translocation of AJ-associated F-actin and β-Catenin-mOrange2, and the predominantly apical extrusion of a central, apoptotic cell. Time-lapse imaging revealed four successive stages defined by the degree of junctional constriction and the apical-basal position of F-actin and β-Catenin (Fig. 2, 7).

**Fig. 7.**
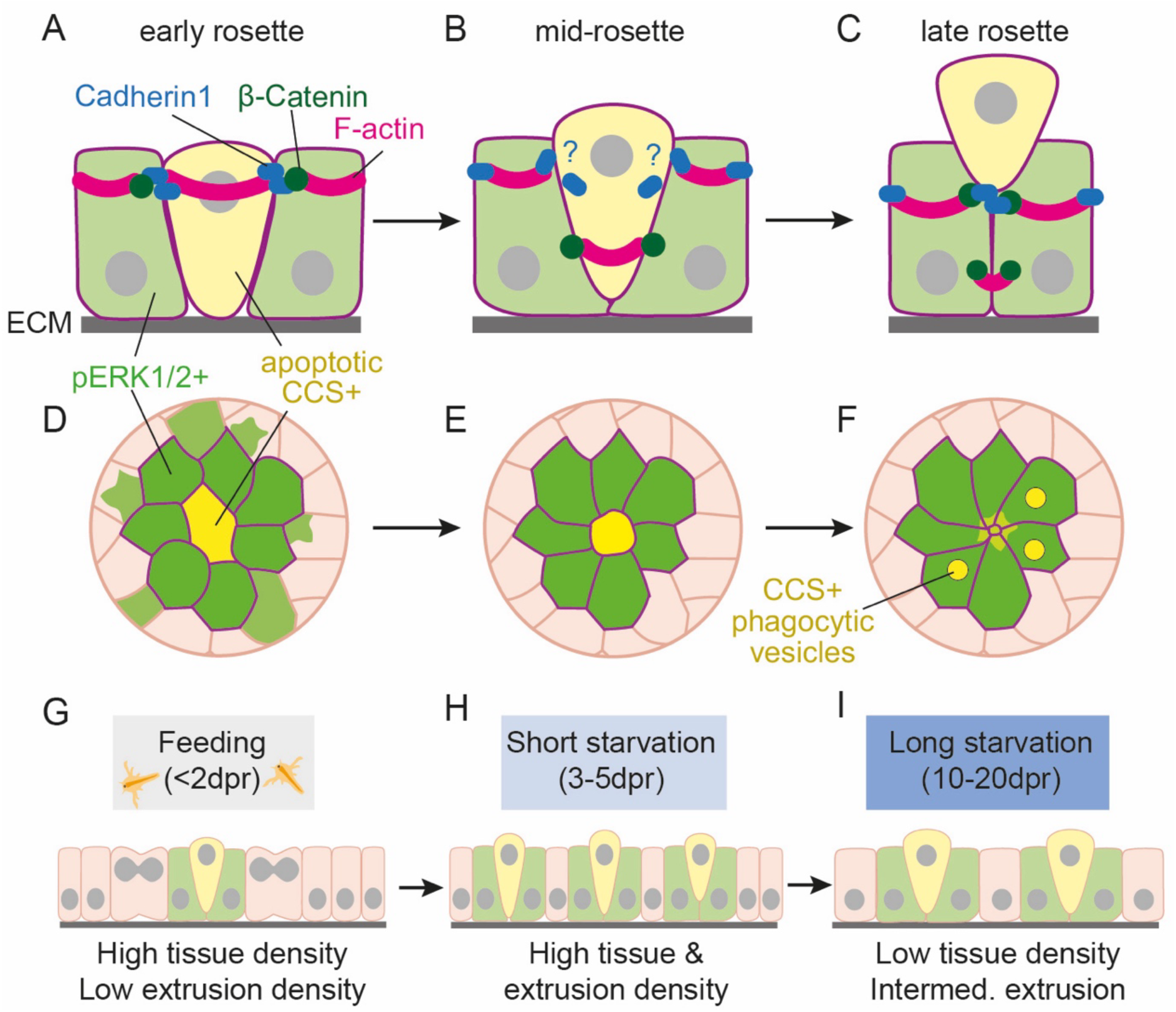
Models of cellular and tissue changes associated with epidermal cell extrusion in *Nematostella*. (A-G) Schematics highlighting changes in adherens junction components (A-C; cross-section), ERK1/2 and Caspase signalling-associated changes (D-F; apical view), and epithelial dynamics associated with cell extrusion (G-I; cross-section) in the body wall epidermis of *Nematostella*. (A-C) During extrusion, F-actin and β-Catenin translocate from their apical position to form a basal, constricting ring, likely providing force for apical extrusion. It remains unclear whether their translocation is specific to the extruding cell or also occurs also in surrounding rosette cells, and whether Cadherin1 proteins likewise translocate extrusion. (D-F) ERK signalling is highest within rosettes but is active at lower levels beyond. The extruding cell undergoes apoptosis as indicated by cleaved Caspase substrate (CCS) immunolabelling, nuclear fragmentation and p38 signalling. Late rosettes occasionally contain CCS+ phagocytic vesicles and active TOR signalling, indicating nutrient recycling by efferocytosis (F). (G-I) Schematic model of the dynamic changes in tissue and rosette density change after a feeding and during starvation. A 1-hour feeding pulse triggers cell proliferation within 48 hours (G), leading to tissue crowding and increased extrusion density (G, H). As starvation progresses, the tissue density decreases, but rosette density remains at intermediate levels indicating continuous, starvation-induced cell loss (I).

In vertebrates and flies, a cell-autonomous contractile F-actin ring in the extruding cell, and a supracellular F-actin ring undergoing apical-basal translocation in surrounding cells are considered hallmarks of the extrusion process (Rosenblatt et al. 2001; Gagliardi and Primo 2019; Staneva and Levayer 2023). However, comparisons between *Drosophila* and different vertebrate systems revealed a substantial diversity in the timing, direction and coordination of ring contraction and AJ remodelling (Rosenblatt et al. 2001; Slattum et al. 2009; Eisenhoffer et al. 2012; Teng et al. 2016; Thomas et al. 2020; Staneva and Levayer 2023; Grata and Levayer 2025). In *Nematostella,* we cannot resolve with our current tools whether the translocating F-actin and β-Catenin rings reside primarily in the central cell, in neighbouring rosette cells or in both. Nonetheless, our data suggest that translocating AJs may actively contribute to extrusion: the basal shift of both F-actin and β-catenin during constriction, together with dynamic interactions involving Cadherin-1, may maintain the epithelial seal while pushing the extruding cell apically (Fig. 7). Extrusion directionality also shows substantial variation across animals, developmental stages and tissues (Grata and Levayer 2025). Vertebrate epithelia generally shed cells apically (Rosenblatt et al. 2001; Gu et al. 2011), whereas delamination in the *Drosophila* larval epidermis typically occurs basally (Nanavati et al. 2020). However, exceptions such as the apical extrusion in the fly gut (Martin et al. 2018) and bidirectional extrusion in the *Xenopus* embryonic ectoderm (Ventrella et al. 2023) highlight that the direction of extrusion is also context-dependent. In *Nematostella,* we predominantly observed apical extrusion events, although basal or lateral extrusion may also occur but be more difficult to detect in our imaging setup.

Another molecular hallmark of extrusion in flies and vertebrates is the wave-like activation of the ERK/MAPK pathway in surrounding rosette cells. During extrusion, ERK1/2 signalling coordinates the spatial distribution of rosettes and provides a pro-survival signal that protects the surrounding epithelium from pro-apoptotic signals emitted by the extruding cell (Gagliardi et al. 2021; Aikin et al. 2020; Takeuchi et al. 2020). In *Nematostella*, we observed consistently high levels of pERK1/2 in surrounding rosette cells from early stages through extrusion, whereas pERK remained undetectable in central cells. In some cases, weaker pERK1/2 signal extended beyond the rosette into adjacent cells. Whether this variable spatial extent reflects different phases of a potential wave-like ERK1/2 signalling pattern, as described in flies and vertebrates (Gagliardi et al. 2021; Aikin et al. 2020; Takeuchi et al. 2020), will require future *in vivo* imaging studies using ERK activity sensors (Harvey et al. 2008; De La Cova et al. 2017; Hirashima 2022). Pharmacological inhibition of ERK phosphorylation with the MEK inhibitor U0126 increased rosette density, supporting a role for ERK in spatially restricting extrusion events. However, because our analyses rely on single time-point measurements, we cannot exclude the possibility that MEK inhibition prolonged the duration of extrusion events, which would also increase apparent rosette density. Developing a segmentation and tracking pipeline for time-lapse imaging, similar to those developed for *Drosophila* pupal notum or wing disc (Valon et al. 2021; Villars et al. 2023), would enable quantitative analysis of extrusion dynamics and help resolve this question.

In vertebrates and flies, the extruding cell is consistently undergoing apoptosis, but the causal relationship between the Caspase-mediated apoptosis initiation and cell extrusion initiation remains debated (Gibson and Perrimon 2005; Shen and Dahmann 2005; Eisenhoffer et al. 2012; Gagliardi and Primo 2019; Valon and Levayer 2019; Grata and Levayer 2025). For example, effector Caspase activation systematically precedes and is required for every cell extrusion in the *Drosophila* pupal notum (Levayer et al. 2016; Moreno et al. 2019), whereas in *C. elegans*, extrusion can occur independently of Caspases (Denning et al. 2012). In *Nematostella*, we show that the central rosette cells consistently exhibit condensed or fragmented nuclei, activated p38/MAPK signalling and high levels of cleaved Caspase substrate (CCS) immunolabelling. Phospho-p38 has previously been associated with apoptotic cells in *Hydra*, planarians and mice (Tursch et al. 2022; Arnold et al. 2016; Hong et al. 2020), and CCS likewise marks apoptotic cells in human cancer cell lines and in *Nematostella* (Xu et al. 2019; Ma et al. 2019; Stokes et al. 2016; Mazloumi Gavgani et al. 2025). Because CCS+ cells appear already during early, and possibly even pre-rosette stages, Caspases-mediated apoptosis in *Nematostella* is likely triggered before extrusion initiates, similar to observations in the *Drosophila* pupal notum (Levayer et al. 2016). This timing suggests that anoikis, where loss of basal membrane attachment induces apoptosis (Frisch and Francis 1994; Frisch 2021), is unlikely to play a major role during epidermal extrusion in *Nematostella*. Experimentally inducing apoptosis with Staurosporine did not alter the density of extrusions, indicating that apoptosis alone is not sufficient to trigger cell extrusion. This contrasts with mammalian cell lines, where apoptosis induction increases the frequency of extrusion (Rosenblatt et al. 2001; Andrade and Rosenblatt 2011). Conversely, inhibiting Caspases in *Nematostella* using Q-VD-OPh decreased extrusion density, consistent with findings in fly and mammalian systems where Caspase inhibition reduces or prevents extrusion (Marinari et al. 2012; Andrade and Rosenblatt 2011). Altogether, our results indicate that Caspase-mediated apoptosis may be necessary, but not sufficient, for extrusion initiation in the epidermis.

We conclude that epidermal cell extrusion in *Nematostella* shares key molecular and morphological hallmarks with extrusion in *Drosophila* and vertebrates, including rosette formation, translocation of AJ components, active ERK signalling in rosette cells, and apoptosis induction in the extruding cell (Rosenblatt et al. 2001; Gagliardi et al. 2021; Valon et al. 2021; Villars and Levayer 2022; Staneva and Levayer 2023; Grata and Levayer 2025). These similarities strongly support the view that epithelial cell extrusion represents an evolutionary conserved mechanism of epithelial remodelling, dating back at least to the last common ancestor of cnidarians and bilaterians. This possibility is consistent with previously described forms of epithelial cell shedding in other cnidarians, such as the continuous apical shedding of cnidocytes at the tentacle tips in *Hydra* (Beckmann and Özbek 2012; Karabulut et al. 2022), or the shedding of symbiont-containing gastrodermal cells during coral bleaching (Gates et al. 1992; Huang et al. 1998; Fang et al. 1997; Sawyer and Muscatine, n.d.; Weis 2008). Similar epithelial cell loss was also observed in other non-bilaterian taxa: in sponges, apoptotic cells are expulsed during metamorphosis (Sogabe et al. 2016) and from the adult choanocyte chamber (De Goeij et al. 2009; Melnikov et al. 2022), and in the placozoan *Trichoplax adhaerens*, apoptotic cell shedding occurs following radiation stress (Fortunato et al. 2021). However, because these processes have not yet been examined at the molecular and cellular levels, it remains unclear whether any of these examples represent *bona fide* extrusion of reflect distinct modes of epithelial cell removal.

Although extrusion has been studied primarily in the contexts of development and disease, its role in food-dependent tissue or body plasticity remains poorly understood. Starvation-induced intestinal shrinkage has been documented in snakes, mice and *Drosophila*, but a dependence on extrusion-driven cell loss has only been demonstrated for *Drosophila* midgut shrinkage (O’Brien et al. 2011; Martin et al. 2018; Secor 2008; McCue 2010; 2012). We therefore developed a machine-learning-based pipeline for computational cell segmentation and rosette identification to determine whether extrusion dynamics in *Nematostella* are compatible with a role in starvation-induced cell loss. We found that epidermal extrusion rates are lowest during continuous feeding, peak after four days of starvation, and then decline to intermediate levels during prolonged starvation. These dynamics mirror previous findings showing that cell loss is highest between two and five days of starvation (Garschall et al. 2024). Notably, extrusion rates are not lowest during prolonged starvation, even though energy levels are expected to be lowest at this stage. Instead, extrusion peaks when cell density is high due to the preceding feeding-induced proliferation burst. Conversely, extrusion remains very low in continuously fed animals despite high proliferation rates, suggesting that short-term starvation may limit extracellular matrix production, thereby creating crowding that promotes extrusion. Together, these observations suggest that extrusion during short-term starvation is primarily density-driven, whereas extrusion during long-term starvation is more directly regulated by nutrient status.

The extensive loss of cells – and thus nutrients – caused directly or indirectly by starvation could be partially mitigated by nutrient recycling in the remaining cells. Indeed, we found cell debris, including nuclear fragments derived from CSS+ apoptotic cells, enclosed in ∼1µm vesicles within rosette cells. This is consistent with efferocytosis-mediated recycling of extruded apoptotic material (Boada-Romero et al. 2020; Torres et al. 2017). Moreover, after prolonged starvation, epidermal TOR signalling became restricted to late-stage rosette cells that frequently contained apoptotic cell debris. These findings independently suggest that nutrient recycling occurs within rosette cells and subsequently activates TOR signalling during extended starvation. Similar mechanisms have been described in planarians, where extruded epidermal cells are engulfed and digested by intestinal phagocytes as part of ongoing tissue turnover (Lee et al. 2024; Lindsay-Mosher et al. 2024). Also in the cnidarian *Hydra*, starvation-induced cell loss is accompanied by efferocytosis of Caspase-3+ apoptotic cells into epidermal cells (Bosch and David 1984; Cikala et al. 1999; Buzgariu et al. 2008).

Our observation that starvation directly or indirectly regulates cell extrusion in sea anemones suggests that extrusion may not only have evolved to maintain epithelial homeostasis, but also to modulate epithelial density and enable nutrient recycling during periods of limited food availability. Notably, placozoans, sponges, and ctenophores also undergo starvation-induced shrinkage, raising the possibility that epithelial extrusion similarly contributes to cell loss and nutrient recycling in these lineages (Granhag and Hosia 2015; Jaspers et al. 2015; Soto-Angel and Burkhardt 2024; Romanova et al. 2022; Maldonado and Young 1999). Future studies will be needed to test whether extrusion represents not only a core mechanism of epithelial remodelling but also an evolutionarily ancient strategy for food-dependent tissue plasticity across animals.

## Methods

### *Nematostella* breeding and husbandry

*Nematostella vectensis* specimens originate from CH6 females and CH2 males collected at Rhode River, MA (Hand and Uhlinger 1992). Adult males and females are kept separately in the dark at 18°C in ∼16ppt filtered seawater (*Nematostella* medium, NM) and fed with fresh *Artemia nauplii* five times per week, with a daily partial water exchange and a monthly full cleaning of the culture boxes. Spawning is induced every three weeks triggered by temperature increase (25°C) and a 12-hour light cycle overnight (Carvalho et al. 2025; Fritzenwanker and Technau 2002). Embryos are maintained at 25°C, and feeding with *Artemia nauplii* begins approximately 7–10 days post-fertilization. Sex was not determined in juveniles; therefore, all data shown include both males and females. All experiments were performed on juvenile polyps of ∼5 mm in length, approximately 1 month after fertilisation

### Immunofluorescence

Juvenile polyps were relaxed in 0.1 M MgCl₂ in *Nematostella* Medium (NM) and then transferred to a plate containing 3.7% formaldehyde/0.5% DMSO in NM (“killing solution”). They were then carefully transferred into a fixation solution of 1× PBS/3.7% formaldehyde/0.5% DMSO/0.1% Tween-20 and fixed overnight at 4°C. Following fixation, samples were thoroughly washed with 1× PBS/0.1% Tween-20 and stored at 4°C for a maximum of 10 days. Samples were then blocked in 1× PBS/5% normal goat serum (NGS)/1% bovine serum albumin (BSA)/0.2% Triton X-100 for two hours at room temperature. Primary antibody incubation was performed in 5% NGS/1% BSA/0.2% Triton X-100 overnight at 4°C. Primary antibody dilutions were used as follows: 1:200 for Cleaved Caspase, 1:50 for phospho-p38 conjugate, 1:200 for phospho-S6 ribosomal, 1:200 for phosphor-ERK1/2, 1:200 for phosphor-p38 and 1:500 for Cadherin1 (see Table 1 for commercial catalogue numbers). Following overnight incubation, tissues were washed in 1× PBS/0.2% Triton X-100 and then blocked in 1× PBS/5% NGS/0.2% Triton X-100 for 30 minutes at room temperature. Hoechst nuclear staining (1:10000, dilution) (ThermoFisher) and Phalloidin F-actin staining (1:50, dilution) and secondary antibodies (1:1000, dilution) (see Table 1 for commercial catalogue numbers) incubated in 1× PBS/5% NGS/0.2% Triton X-100 overnight at 4°C. Finally, samples were thoroughly washed in 1× PBS/0.2% Triton X-100, cleared in glycerol, and mounted on slides (Electron Microscopy Sciences, 63418-11) in 80% glycerol. Coverslips (Menzel-Gläser, 18×18 mm) were sealed with clear nail polish. Exceptionally, rabbit anti-p-RPS6 was blocked for three nights (Pascual-Carreras et al. 2025). For the cadherin antibody, tissues were instead fixed for 1 hour at 4°C with Lavdovsky’s fixative (3.7% formaldehyde, 50% ethanol, 4% acetic acid) (Pukhlyakova et al. 2019). For double immunolabelling, polyps were incubated with phospho-p38 Alexa Fluor 594 conjugate at a 1:50 dilution along with the phospho-ERK1/2 antibody at a 1:100 dilution in blocking buffer at 4°C overnight. The protocol then proceeded as usual, except that no secondary antibody was necessary for the phospho-p38 Alexa Fluor 594 conjugate (Tursch et al. 2022). Tissue staining and immunohistochemistry has been performed on different feeding states including early post-feeding (1-2, 5 days starved), as well as late starvation (20 days starved). In the manuscript exemplary images represent the results found across multiple feeding states unless a significant different result was found between fed and starved states.

**Table 1:** Key resources table.

| REAGENT or RESOURCE | SOURCE | IDENTIFIER |
| --- | --- | --- |
| **Antibodies** |  |  |
| Cleaved Caspase Substrate Motif [DE(T/S/A)D] MultiMab® | Cell Signaling Technology | Cat# 8698 |
| Phospho-p38 MAPK (Thr180/Tyr182) (D3F9) XP® (Alexa Fluor® 594 Conjugate) | Cell Signaling Technology | Cat# 8632S |
| Anti-Phospho-S6 Ribosomal Protein (Ser235/236) | Cell Signaling Technology | Cat# 4858 |
| Phospho-p44/42 MAPK (Erk1/2) (Thr202/Tyr204) (D13.14.4E) XP® | Cell Signaling Technology | Cat# 4370T |
| Phospho-p38 MAPK (Thr180/Tyr182) (D3F9) XP® | Cell Signaling Technology | Cat# 4511T |
| Nv_Cadherin1 | (Pukhlyakova et al. 2019) | N/A |
| Goat anti-Rabbit IgG (H+L) DyLight 488 conjugate | LifeTech | Cat# 35552 |
| Goat anti-Rat IgG (H+L), Alexa Fluor® 633 conjugate | LifeTech | Cat# A21094 |
| AZD8055 | Nordic Biosite | Cat# hy-10422 |
| Staurosporine | Sigma Aldrich | Cat# S4400 |
| U0126 | Merck | Cat# 662005 |
| QV-D-OPh | MedChemExpress | Cat# hy-12305 |
| TrueCut Cas9 protein v2 | ThermoFisher | A36498 |
| **Deposited data** |  |  |
| *Nematostella vectensis* genome | (Putnam et al. 2007) | https://mycocosm.jgi.doe.gov/Nemve1/Nemve1.home.html |
| **Experimental models: Organisms/strains** |  |  |
| *Nematostella vectensis* | Michael Sars Centre | N/A |
| *Nematostella*: β-Catenin-mOrange2 fusion protein transgenic knock-in line | This study | N/A |
| **Oligonucleotides** |  |  |
| Primers for gRNA oligos and primers for PCR and sequencing of the transgenic line can be found in Table 2 | This study | N/A |
| Recombinant DNA |  |  |
| Plasmid: β-Catenin-mOrange2 (pJet1.2 backbone) | This study | N/A |
| Software and algorithms |  |  |
| Primer3Plus | (Untergasser et al. 2012) | http://www.bioinformatics.nl/cgi-bin/primer3plus/primer3plus.cgi |
| CRISPOR | (Concordet and Haeussler 2018) | http://crispor.tefor.net/ |
| Fiji | (Schindelin et al. 2012) | https://imagej.net/software/fiji/ |
| Cellpose | (Stringer et al. 2021)(Stringer and Pachitariu 2025; Pachitariu and Stringer 2022) | https://www.cellpose.org |
| Napari | (Chiu et al. 2022) | https://napari.org/dev/index.html# |
| R Studio | (R Core Team 2025) | v.4.5.2<br>https://www.r-project.org/ |

### Drug treatments

All pharmacological inhibitors were prepared from 100% stocks and diluted in NM to their working concentrations: AZD-8055 from 10mM stock to 0.1µM (Pascual-Carreras et al. 2025), Staurosporine from 500µM to 0.5 μM; U0126 from 10mM to 20 μM, and QV-D-OPh from 10mM to 100 μM. All treatments had additional DMSO added to a final concentration of 0.5% in NM, and the negative controls included 0.5% DMSO to match treatments (see Table 1 for commercial catalogue numbers), except for QV-D-OPh which was used at 1% DMSO for live imaging. For all drug treatments, polyps were placed in drug solution for 2 hours at 25°C standard culturing temperature before live imaging. 30 min before drug removal, some drops of 1M MgCl_2_ were added to relax animals and soon transferred into fresh NM with 0.1M MgCl_2_ followed by mounting into in Ibidi µ-Slide VI 0.4 (Cat. # 80606) and confocal imaging.

### Transmitted light and confocal imaging

Immunofluorescence samples were imaged using an Olympus FV3000 confocal microscope (standard PMT detectors, 63× silicon-immersion objective). Live juvenile polyps were relaxed in 0.1 M MgCl₂ in NM, mounted on an Ibidi µ-Slide VI 0.4 (Cat. # 80606) and also imaged on the Olympus FV3000 confocal microscope. Confocal room temperature was at a range between 20-22 °C and therefore live imaging was performed at that temperature. Images and confocal stacks were processed, cropped, and adjusted for levels and colour balance using ImageJ/Fiji.

### CRISPR/Cas9-mediated generation of β-Catenin-mOrange2 knock-in line

The used protocols were adapted from previously established protocols for the generation of CRISPR/Cas9-mediated knock-in lines in *Nematostella vectensis* (Lebouvier et al. 2022; Miramón-Puértolas et al. 2024). A single guide RNA (gRNA) was designed with a predicted cut site 14bp upstream of the Stop codon of the *Nematostella* β-catenin gene (Nvec200_v1.10389.1.p1) using CRISPOR (Concordet and Haeussler 2018). The gRNA template was generated by annealing and elongating an invariant T7 promotor/tracrRNA-encoding and a sequence-specific oligos (Bassett et al. 2013) (ThermoFisher, desalted, see Table 2). *In vitro* transcription of the gRNAs was performed using a T7 MegaScript Transcription Kit (ThermoFisher), followed by ammonium chloride precipitation and dilution in nuclease-free water to a final concentration of 1.5 μg/μl. The donor DNA fragment for homology-mediated repair (HDR) consisted of the 3’-end of the *β-catenin* gene open reading frame (ORF) cloned in-frame with a (GGGGS)2 linker and the mOrange2 protein-coding sequence. The construct also contained 994 bp long ‘left’ (upstream) and 936 bp long ‘right’ (downstream) homology arms. This entire construct was cloned into a pJet1.2 plasmid backbone (ThermoFisher) using Gibson Assembly Master Mix (NEB) (Gibson et al. 2009). The donor DNA fragment was PCR-amplified using biotin-labelled oligos (ThermoFisher; desalted, see Table 2) flanking the donor fragment to enhance integration efficiency (Gutierrez-Triana et al. 2018). Cas9-mediated knock-in injections were carried out by modifying existing protocols for CRISPR/Cas9-mediated mutagenesis, using 0.75 μg/μl TrueCut Cas9 protein v.2 (Thermofisher), 75 ng/μl of each guide RNA, 70 ng/μl column-purified donor DNA, and a modified injection buffer containing 220 mM KCl (Kraus et al. 2016; Burger et al. 2016). Approximately 1600 zygotes were injected, and the resulting juvenile polyps displaying mOr2+ signal were selected and raised to sexual maturity. At the adult stage, polyps exhibiting mOrange2 signal in the developing gametes were chosen as founder animals and outcrossed with wild-type animals to generate the F1 generation. The successful integration of the *linker-mOrange2* fragment knock-in was validated at the transcript level by cDNA synthesis from total RNA extracted from a mix of larvae tissue resulting from a cross of two heterozygous F1 animals. Using this cDNA, regions spanning the transition between the 3’ UTR and mOrange2-encoding region, and between the mOrange2-encoding region and the 3’ end of the *β-catenin* ORF were amplified using oligos outside the homology arm regions (see Table 2). Sanger sequencing of PCR fragments flowing gel elution and column purification confirmed the flawless, in-frame integration of the *linker-mOrange2* fragment into the locus. The successful, flawless genomic integration was further validated at the genomic level by extracting genomic DNA from mixed tentacle clips of several homozygous F2 animals from the same founder F1 animals. Based on this genomic DNA, the transitions between the 3’-end of the *β-catenin* ORF and the (GGGGS)2 linker/mOr2 and between and the 3’ end of the mOr2 ORF and the 3’UTR of the *β-catenin* gene were amplified using oligos outside the homology arm regions (see Table 2). Gel elution and Sanger sequencing confirmed successful integration.

**Table 2:**
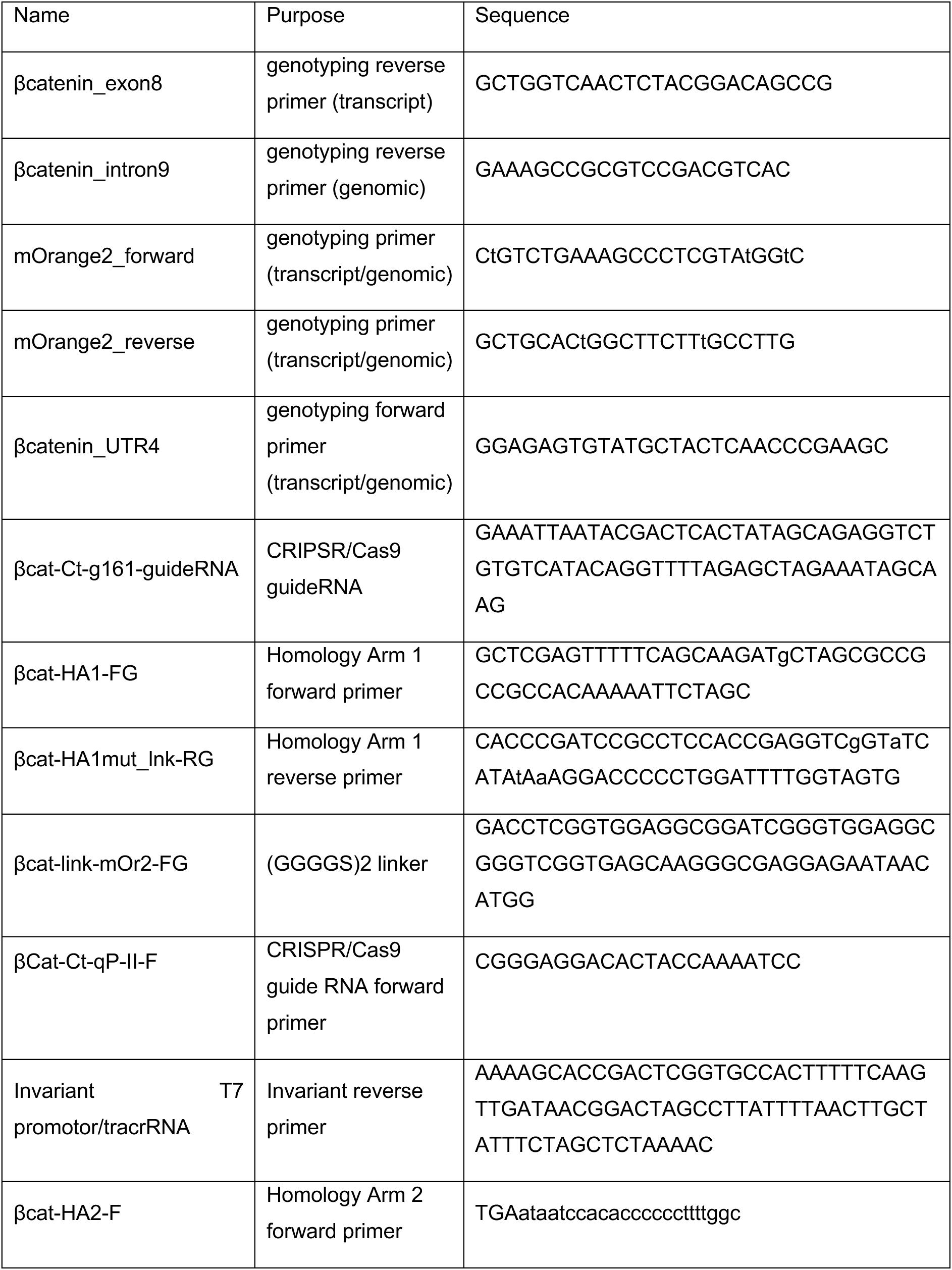
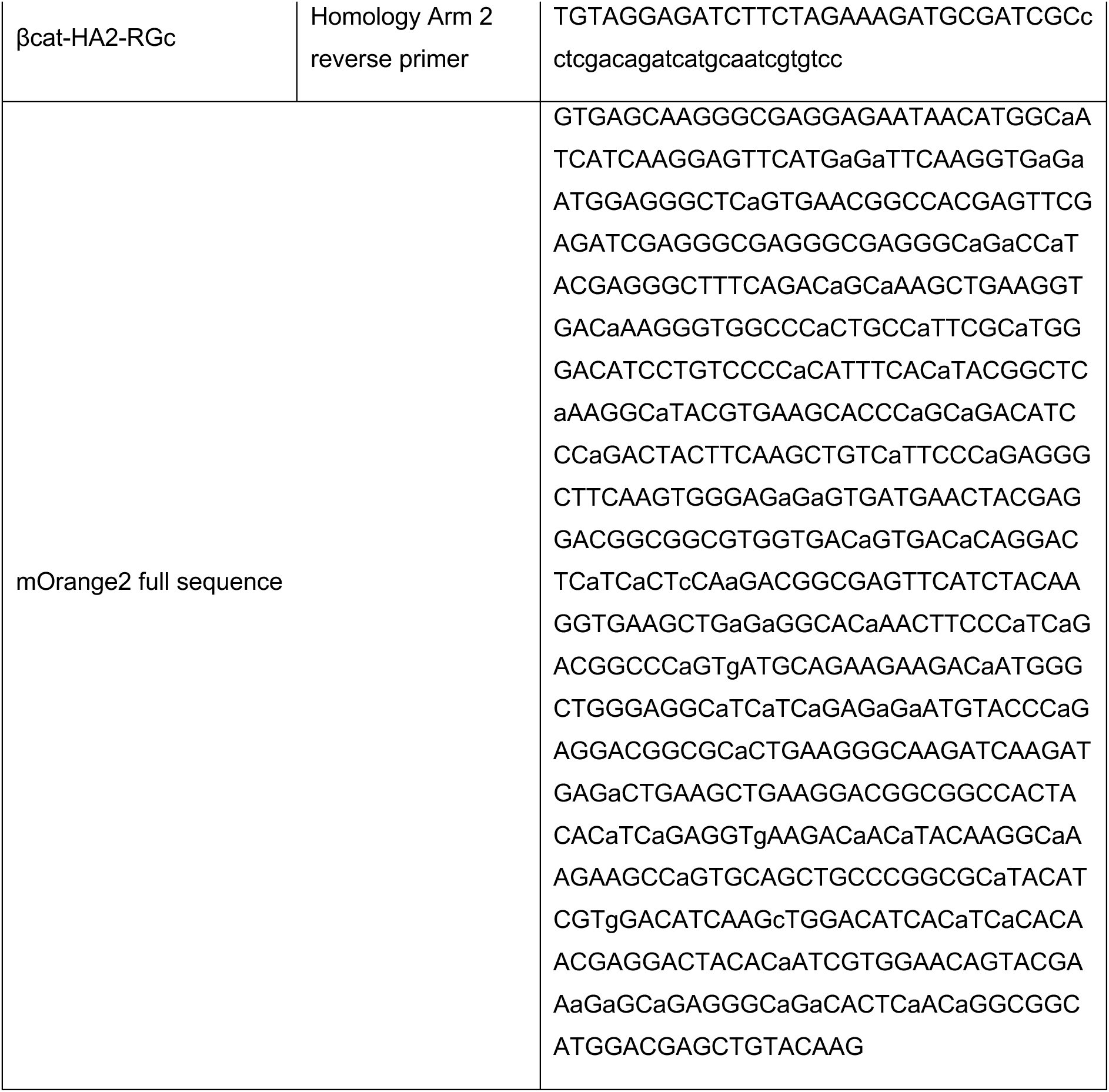
List of single guide RNAs, corresponding melt-curve analysis primers and mutant genotyping primers for the generation of β-catenin-mOrange4 knock-in line.

### Image acquisition and experimental design

Data acquisition was performed using a confocal microscope. For each β-catenin-mOrange2 transgenic juvenile *Nematostella vectensis* polyp, two to three sets of image stacks were acquired, focusing on the epidermal cell layer. Approximately 10-15 animals per treatment condition or timepoint were imaged to ensure sufficient statistical power. In the starvation experiment, animals were initially starved for five days to control for their feeding history (T_5ds_). They were then refed for 1 hour and subsequently sampled at the following timepoints: 1 day post-refeeding (dpr) through to 5 dpr, 10 dpr, and 20 dpr. At 20 dpr timepoint, animals were fed again for 1 hour and sampled twenty-four hours later to obtain a final timepoint (+24h). This starvation experiment was conducted with two biological replicates.

For the drug treatments, animals were starved for 5 days (T_5ds_) fed for 1 hour and then starved for either 5 dpr or 20 dpr before being treated with one of four compounds: AZD8055, U0126, QV-D-OPh, or Staurosporine (see details above for Drug treatments). Each treatment condition had 3 biological replicates and corresponding DMSO controls. In total, this resulted in two timepoints (5 and 20 dpr), four inhibitors, four respective controls, and three independent replicates per condition, a total of 937 images. As with the starvation data, two to three image stacks were acquired per animal, with approximately 10-15 animals imaged per treatment or control group.

Juvenile polyps used for the daily fed dataset were maintained under the normal daily feeding regime used for general *Nematostella vectensis* husbandry. Animals were fed daily each morning and then cleaned in the early afternoon to remove residual food and waste before imaging. Imaging was performed at the same time of the day (14.00-16.00) as all other experiments (starvation and drug) to standardise conditions across all samples. For each of the three consecutive days, a different biological batch of animals was used, providing three independent replicates. This resulted in three separate datasets representing animals under daily feeding conditions. As with the starvation and drug experiments, two to three image stacks were acquired per animal, with approximately 10–15 animals imaged per biological replicate.

### Image curation and cell segmentation pipeline

Following image acquisition, the image stacks were manually curated to select the most appropriate z-planes, specifically those where the surface of the epidermis was best visualised. Image z-stacks were then edited into 2D maximum intensity projections with the use of ImageJ/Fiji (Schindelin et al. 2012) to improve the coverage of the epithelium especially in cases of capturing areas in which the epidermis was not entirely flat while also improving the focus of the image (Brocher 2025). These curated images were then subjected to initial segmentation using Cellpose software (https://www.cellpose.org) (Stringer and Pachitariu 2025; Pachitariu and Stringer 2022). For the time-series dataset, approximately 250 of 500 images were manually corrected after initial segmentation to refine cell boundaries and eliminate segmentation errors. These manually corrected masks were then used to train a custom Cellpose Cyto3 base model, which improved performance and was subsequently used to segment the remainder of the time-series dataset as well as the inhibitor dataset. Post-segmentation processing included computational quality control steps to remove background artefacts, identify unsegmented regions, and classify cells based on size. Segmented objects below a defined pixel area threshold were assumed to be “oversegmented” or background noise and were merged with neighbouring cells. Conversely, objects with abnormally large areas were excluded as background or segmentation artefacts. The resulting processed masks were used for further analysis.

### Rosette annotation and prediction

For automatic rosette segmentation, we trained an Attention U-Net deep learning model (Oktay et al. 2018) on a manually annotated subset of 268 time-series images. To prioritise multicellular geometric features over intensity cues, the model was trained exclusively on binary cell masks and morphologically dilated boundaries rather than raw fluorescence. Model performance was monitored on a 20% held-out validation set using Dice and F1 scores. Following inference, all predictions underwent manual review and correction. We quantified the method’s efficiency by calculating the speed-up factor of this model-assisted workflow compared to two baselines: manual identification on segmented images and manual identification on raw data (see further details in Supp. Methods). To ensure reproducibility and facilitate scalable processing across all datasets, the entire analysis workflow — from segmentation to quantification — was orchestrated using a custom Snakemake pipeline (Köster and Rahmann 2012), with software environments managed via Poetry (Eustace and Contributors 2018) (see further details in Supp. methods).

### Statistical analysis and graphical representation of data

Data visualization and analysis were performed using R (v. 4.5.2 (2025-10-31), https://www.r-project.org/, (R Core Team 2025)) with the packages: mclust (v. 6.1.2 (Scrucca et al. 2023)), ggplot2 (v. 4.0.0 (Wickham 2016)), dplyr (v. 1.1.4 (Wickham et al. 2023)), tidyr (v. 1.3.1 (Wickham et al. 2024)), lme4 (v. 1.1.37 (Bates et al. 2025)), lmerTest (v. 3.1.3 (Kuznetsova et al. 2017), using Type III ANOVA, Satterthwaite’s approximation), and emmeans (v. 2.0.0 (Lenth and Piaskowski 2025)). Cell area measurements exhibited a robust bimodal distribution consisting of two populations of small and large cells. To statistically define these classes, a Gaussian Mixture Model (GMM) with two components was fitted to each image (see Supp. methods for details). Cells were classified as small or large based on Bayesian posterior probabilities. Across all datasets, the proportions of cell classes were stable (∼25% small, ∼75% large), and the size of the small-cell population remained invariant across treatments and timepoints. As a result, small cells were excluded from analyses of rosette density. Rosette density was calculated as the number of rosettes per 1000 large cells per image. For the rosette-density timeseries and daily fed datasets, temporal changes were assessed using linear mixed-effects models to account for replicate variability and to obtain model-adjusted means and p-values. Cell area dynamics were analysed separately for small and large cells. Cell area was also modelled using linear mixed-effects models from which estimated marginal means (EMMs) and Tukey-adjusted pairwise comparisons were extracted. Additional details on mixed-model fitting are provided in supplementary methods. For drug experiments, treated vs. control comparisons were assessed using Wilcoxon rank-sum tests and Kruskal–Wallis tests, supplemented with rank-transformed mixed models (see further details in Supp. methods). All reported means in the main text, p-values, and percentage changes refer to EMM-adjusted results. Rosette density and cell area were visualised using boxplots and violin plots, respectively. Figures were exported as vector graphics (PDF), and final layouts were refined in Adobe Illustrator.

## Acknowledgements

We thank Sara Brites for experimental support, all past and present members of the Steinmetz lab, especially Kathrin Garschall, for support and discussions, Eilen Myrvold, Brandon Mellin and Lavina Jubek for taking excellent care of the *Nematostella* culture at the Michael Sars Centre Cnidarian Facility, Christopher Noack from the Bosch Lab at the University of Kiel for generously sharing Ibidi μ-Slide experience and test samples, and Marios Chatzigeorgiou and Andreas Midlang for their generous help and support of confocal microscopy imaging.

## Author contributions

Conceptualization, I.F.-F., L.C. and P.R.H.S.; Methodology, I.F.-B., N.B. and P.R.H.S.; Validation, I.F.-B.; Investigation, I.F.-B., Formal analysis, I.F.-B. and N.B.; Data curation, I.F.-B. and N. B.; Visualization, I.F.-B. and P.R.H.S.; Writing - Original Draft, I.F.-B. and P.R.H.S.; Writing – Review & Editing, I.F.-B., N.B., L.C. and P.R.H.S.; Resources, P.R.H.S. and L.C.; Funding Acquisition, L.C. and P.R.H.S.; Supervision, L.C. and P.R.H.S.

## Lead contact

Further information and requests for resources and reagents should be directed to and will be fulfilled by the Lead Contact, Patrick Steinmetz.

## Code availability

The complete source code and automated analysis pipeline are available at https://github.com/noahbruderer/nematostella_rosette_detection under the MIT License.

## Declaration of Interests

The authors declare no competing or financial interests.

## Declaration of generative AI and AI-assisted technologies in the manuscript writing process

During the preparation of this manuscript, Large Language Models (OpenAI, https://chat.openai.com, Claude by Anthropic, https://claude.ai/ and Gemini by Google, https://gemini.google.com) were used to assist with coding and R script troubleshooting, as well as for polishing and grammar checking of the manuscript text. All outputs were critically reviewed and edited, and the authors take full responsibility for the final content.

## Funding

All authors were supported by the Michael Sars Centre core budget that received funding from the University of Bergen and the Norges Forskningsråd (234817). https://www.uib.no/en. None of the funders played any role in the study design, data collection and analysis, decision to publish, or preparation of the manuscript.

## Notes

https://github.com/noahbruderer/nematostella_rosette_detection

### Competing Interest Statement

The authors have declared no competing interest.

