## Supplementary_figures for "Epithelial cell extrusion underlies starvation-induced cell loss in a sea anemone"

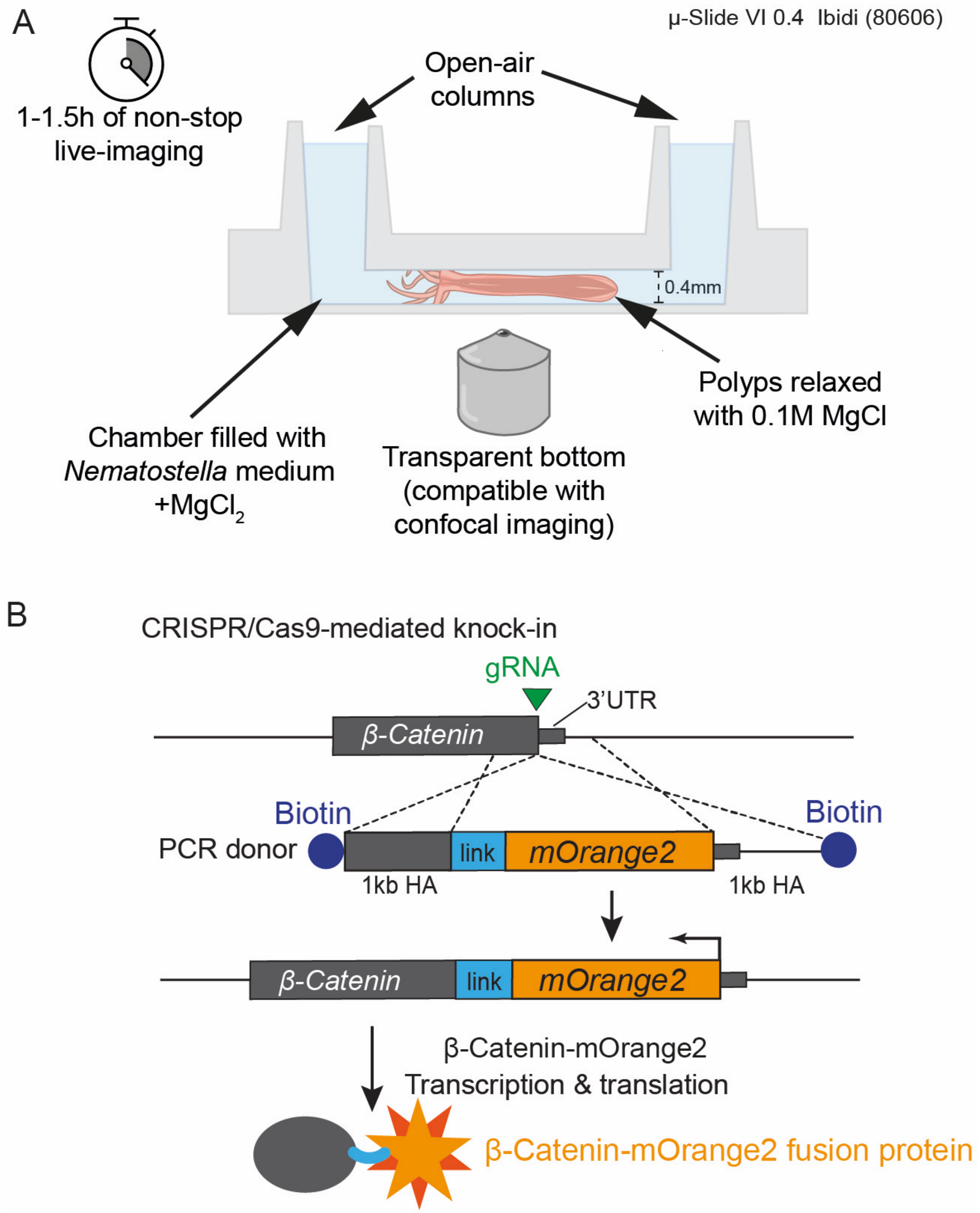


**Fig. S1. Live imaging set up and strategy for generating a β-Catenin-mOrange2 fusion protein transgenic line.**

(A) Live imaging set-up for confocal imaging of *Nematostella* juvenile polyps. Juveniles were immobilized and imaged in commercially available Ibidi μ-Slide VI (80606) chambers with a 0.4mm channel height. Each chamber provides two open ports that allow continuous gas exchange, offering a more stable environment for live polyps compared to sealed coverslip preparations. (B) Schematic depicting the CRISPR/Cas9-mediated homology-directed recombination strategy to generate a β-Catenin-mOrange2 knock-in allele. A single gRNA was used to induce a double-strand break close to the stop codon of the β-Catenin open-reading frame. Homology-directed repair was mediated by a donor construct with ±1 kb homology arms, encoding a linker ($GGGGS_2$) and encoding the mOrange2 reporter protein cloned in-frame to result in a C-terminal fusion protein. The donor DNA was biotinylated at the 5'-ends to prevent concatemerization. Successful, flawless integration was validated on both transcript and genome levels (see Methods). Abbreviations: h: hour, gRNA, guide RNA; HA, homology arm; kb, kilobase; ORF, open reading frame and UTR, untranslated region.

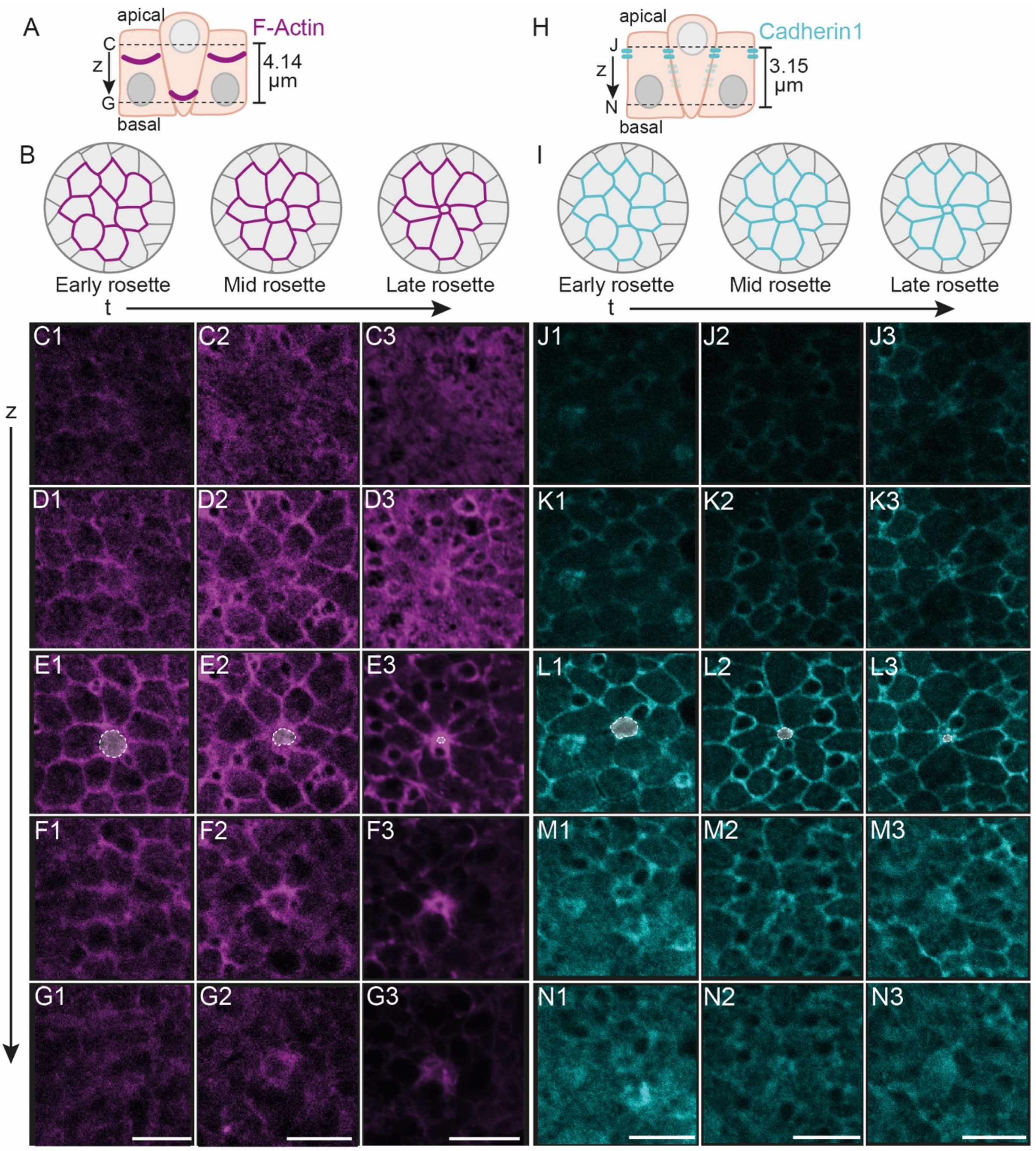


**Fig. S2. Cadherin and F-actin are dynamically distributed along the apical-basal axis during cell extrusion.**

(A, H) Schematics indicating the relative positions of F-actin (A) or Cadherin1 (H) and of image panels (in C1-G3, J1-N3) along the z plane. (B, I) Schematic overviews of typical rosette shapes at early rosette, mid-rosette and late rosette stages. (C1-G3, J1-N3). Single plane confocal microscopy images of fixed juvenile body wall epidermis labelled by Phalloidin (F-actin; C1-G3) or an antibody detecting *Nematostella* Cadherin1 (J1-N3) along the z-axis from apical (C, J) to basal (G, N) and from early rosette (C1-G1, J1-N1) to late rosette stages (C3-G3, J3-N3). Shaded area in E1-E3,

L1-L3 highlight progressive constriction and extrusion of the central cell. Note the broad apical localisation of Cadherin1 and basal translocation of the central F-actin ring during extrusion. Scale bars: 10 μm.

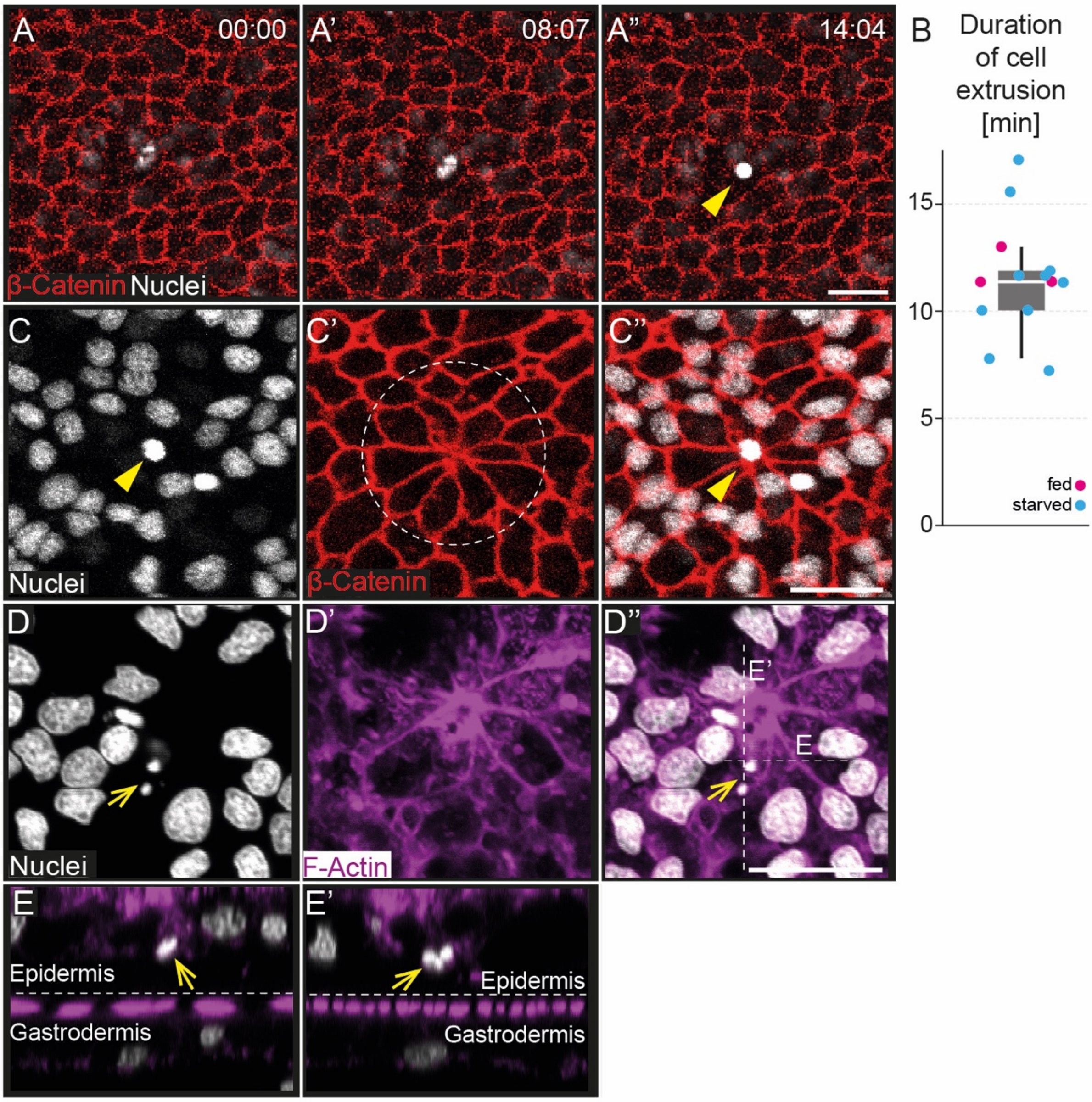


**Fig. S3. Examples of apical and putative basolateral cell extrusion events.**

(A-A'', C-C'') Single plane *in vivo* confocal microscopy images of time-lapse recordings of β-Catenin-mOrange2 transgenic juvenile polyps combined with Hoechst nuclear staining. The condensed nuclei (yellow arrowheads) at the centre of a rosette indicates the apoptotic nature of the extruding cell, and its apical appearance (in A-A'') at the level of AJs indicates apical extrusion (see also Supplementary Video 2). (B) Box plot showing the spread in duration of cell extrusion events based on *in* vivo time-lapse recordings of fed and starved juvenile polyps (n = 13, consisting of 3 fed polyps (1-2 days after feeding) and 10 polyps starved for ca. 20 days). Box plot shows median value (middle bar), the first to third interquartile ranges (box) and whiskers indicate 1.5x IQR. Individual datapoints are plotted as dots, coloured depending on their feeding status (fed: fed; starved for 20 days: blue). (D–E′) Projection of confocal stack

(D-D'') and z-axis reconstructions (E,E') of late rosette in juvenile polyp epidermis stained with phalloidin (F-actin) and nuclear staining (Hoechst) reveals a basally positioned, highly fragmented nucleus (yellow arrows) indicating basal or lateral extrusion. Nuclear stain: Hoechst. Scale bars: 10 μm. Time: min:sec

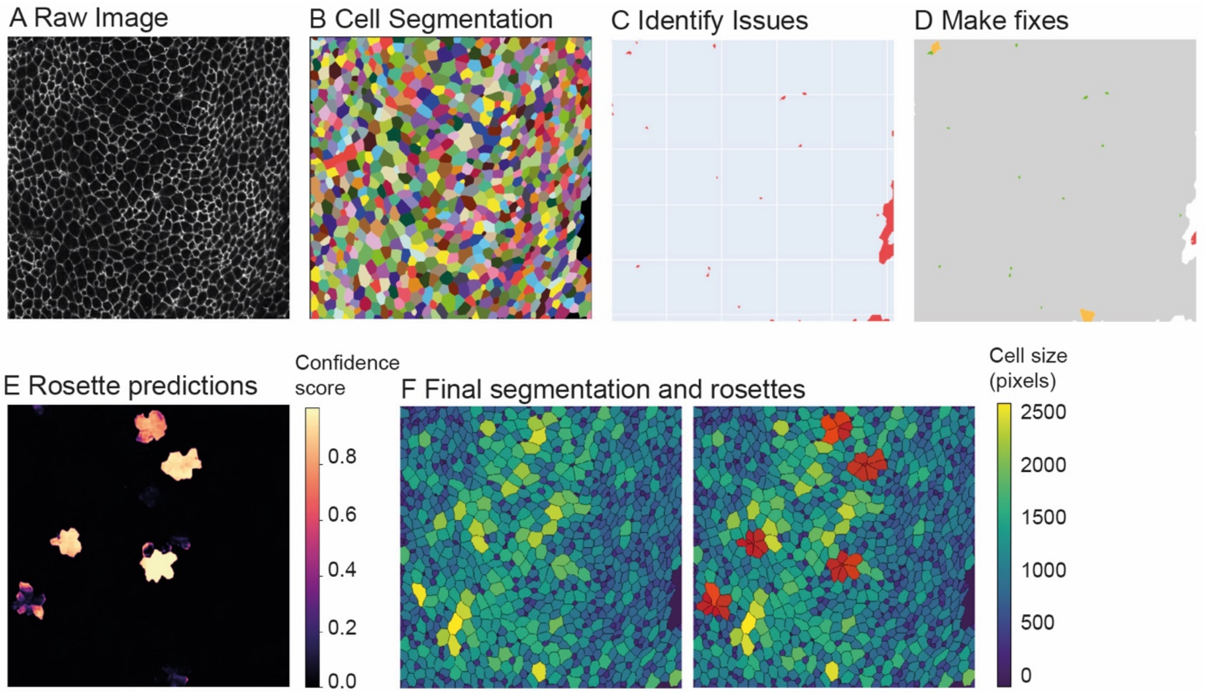


**Fig. S4. Illustrative example images of image segmentation and rosette ID pipeline from the confocal image projection to assigned rosettes and cell outlines**

(A) Projection of *in vivo* confocal microscopy image showing cell outlines labelled using a β-Catenin-mOrange2 transgenic juvenile polyp. Image represents an example used for subsequent image segmentation. (B) Result of initial cell segmentation using Cellpose on the image in (A), generating a pre-processed mask. (C-D) Quality control based on a cell segmentation image (B) to identify (C) and correct (D) for artefacts such as background, unsegmented, very small or very large cells. Cleaned mask after filtering out background, excluding large artefacts, and merging small fragments with neighbours (D). (E) Rosette likelihood map based on manually trained model. (F) Final annotated segmentation combining both cell outlines and rosette assignments.

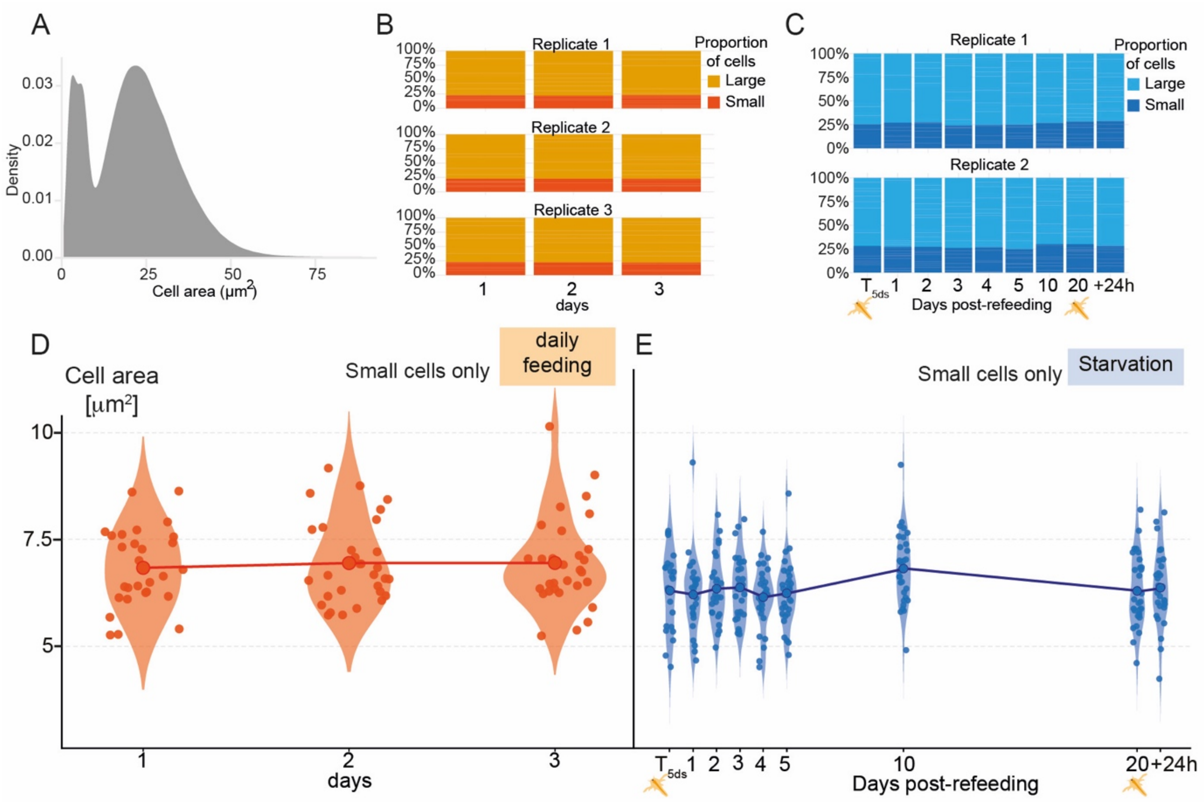


**Fig. S5. Quantification of cell area changes in daily fed and starved juvenile polyps reveals two populations of large and small cells**

(A) Binomial distribution of cell areas in the epidermis of all pooled polyps across the starvation dataset. (B, C) Bar plots show the relatively invariant proportions of cells with small (red, dark blue) and large (orange, light blue) areas per replicate in the 'daily fed' (B) or starvation (C) datasets. (D-E) Violin plots represent the distribution of small cell area during daily feeding (D) or starvation experiments (E; see Fig. 3A, B for setup). Note that the cell area of small cells stays within a narrow range across all sampling timepoints (D, E), while the cell area of larger cells changes dynamically during starvation (see Fig. 3D). Larger central dots represent means per timepoint, which are joined to the other means to form a trend line. Smaller dots represent mean small cell area per animal.

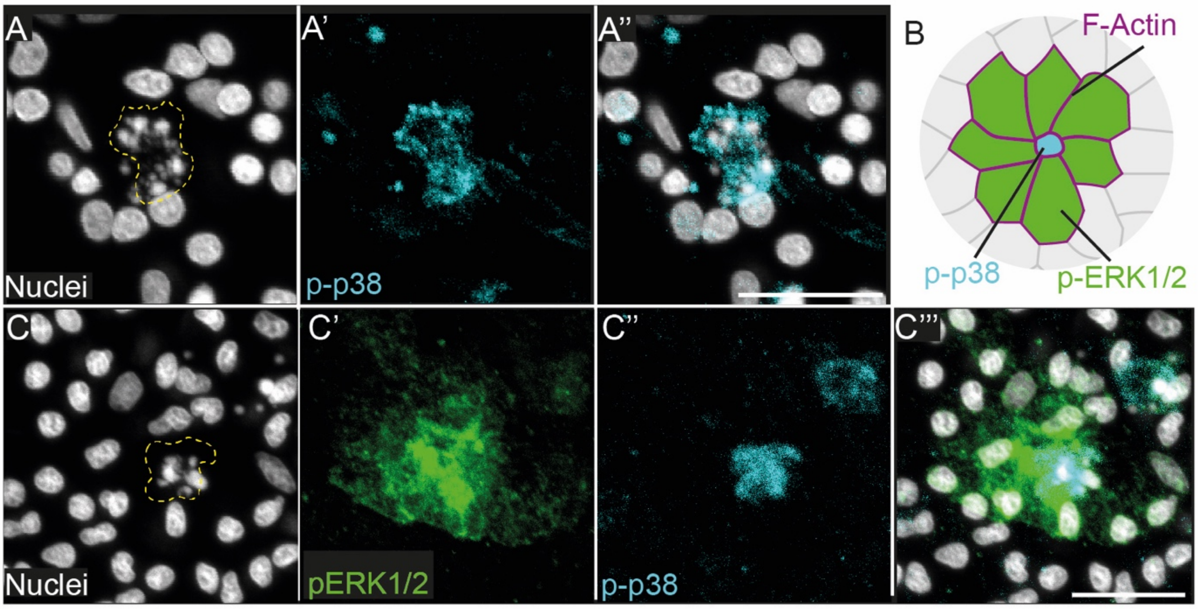


**Fig. S6. Phosphorylated p38-labelled (p-p38+) cells with fragmented nuclei located at the centre of phosphorylated-ERK1/2 (pERK1/2+) rosette.**

(A–A'', C-C''') Projections of confocal imaging stacks of fixed epidermis labelled for p-p38, pERK1/2 and nuclei. Note the nuclear fragmentation (yellow dotted area) as an indication of late apoptosis in the p-p38+ cells within a pERK1/2 patch. Nuclear stain: Hoechst. Scale bars: 10 µm. (B) Schematic summary of pERK1/2 and p-p38 labelling that together support that extruding cells are p-p38+.

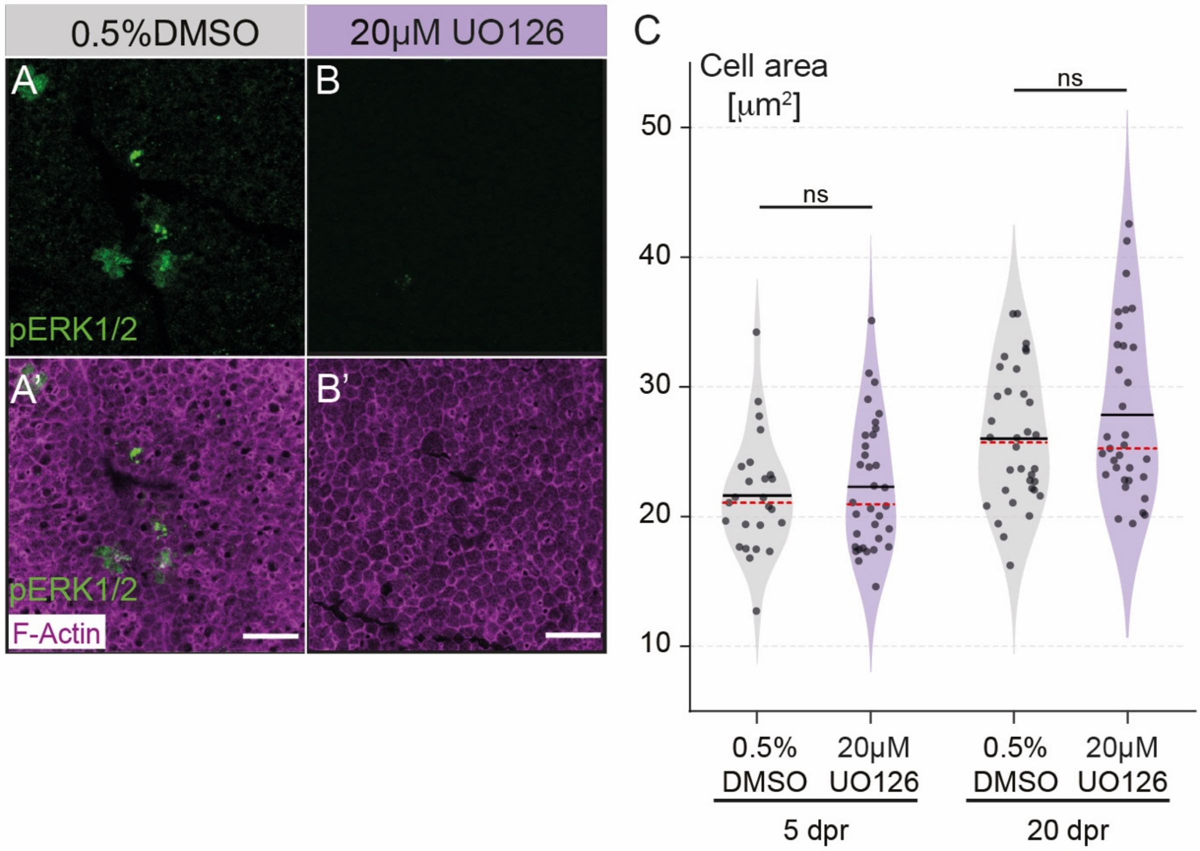


**Fig. S7. MEK inhibition by U0126 suppresses ERK1/2 phosphorylation without significantly affecting the cell area distribution of large cells.**

(A-B') Projections of confocal imaging stacks of fixed epidermis of 5 days-starved polyps labelled for phosphorylated ERK1/2 (pERK1/2) and F-actin (phalloidin) after 2 hours incubation in 0.5% DMSO (A–A') or 20 µM U0126 (B–B'). Images are representative of the whole-body wall epidermis, with pERK1/2+ cell patches becoming undetectable after U0126 incubation. (C) Violin plots show the distribution of mean cell areas across larger cells in the epidermis after 2 hours of 0.5% DMSO (control) or 20 µM U0126 treatments at 5 days post-refeeding (dpr) or 20 dpr. Note that U0126 does not significantly change tissue density. Means are represented by black full lines and medians as red dashed lines. Single data points represent means of large cell areas per animal. Significance codes for model adjusted P-values: ns = non-significant. Abbreviations: dpr: days post-refeeding. Scale bars: 20 µm.

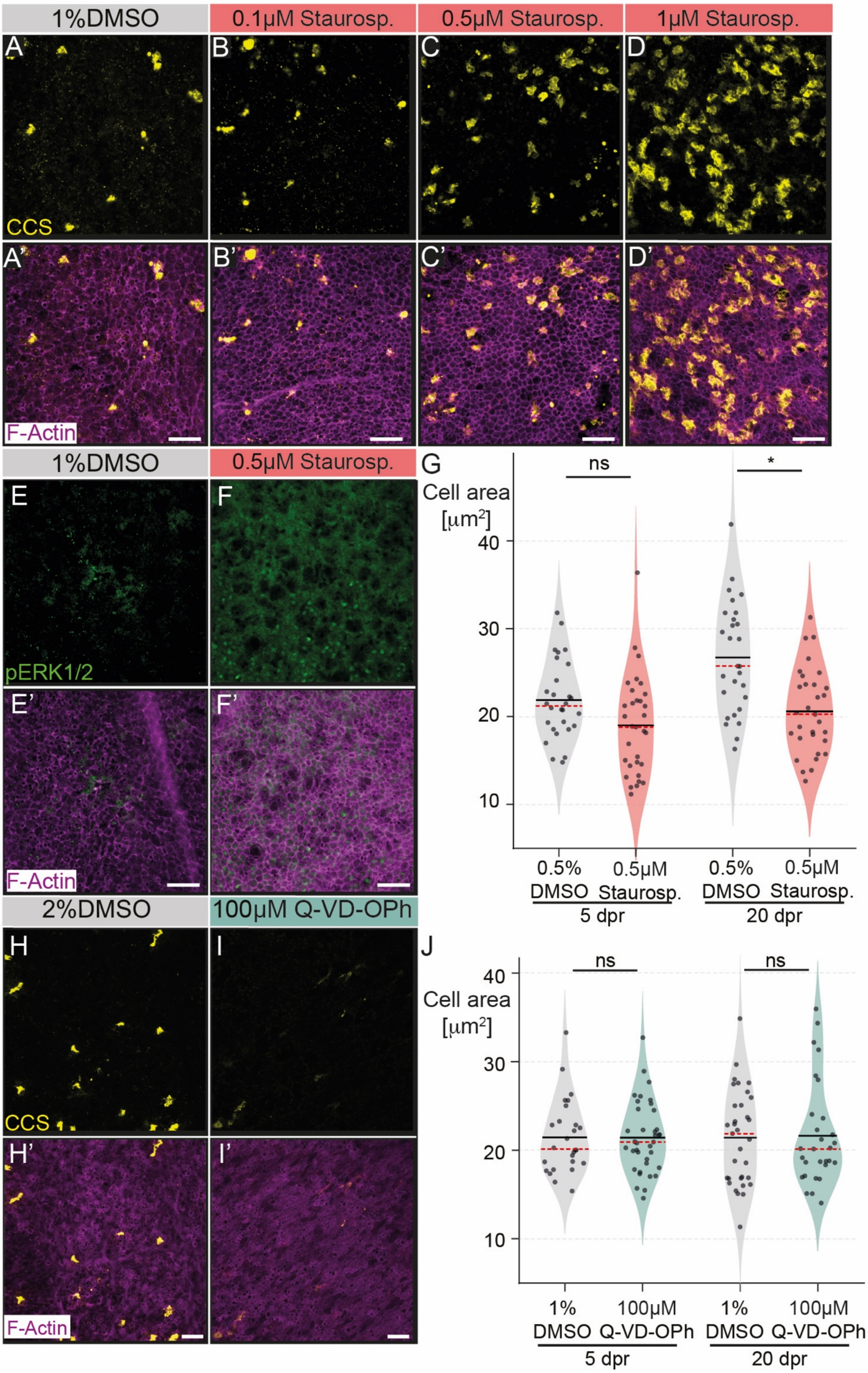

1%DMSO
0.1μM Staurosp.
0.5μM Staurosp.
1μM Staurosp.
A
B
C
D
CCS
A'
B'
C'
D'
F-Actin
1%DMSO
0.5μM Staurosp.
E
F
pERK1/2
E'
F'
F-Actin
G
ns
*
Cell area
[μm2]
40
30
20
10
0.5% DMSO
0.5μM Staurosp.
0.5% DMSO
0.5μM Staurosp.
5 dpr
20 dpr
2%DMSO
100μM Q-VD-OPh
H
I
CCS
H'
I'
F-Actin
J
ns
ns
Cell area
[μm2]
40
30
20
10
1% DMSO
100μM Q-VD-OPh
1% DMSO
100μM Q-VD-OPh
5 dpr
20 dpr

**Fig. S8. Effects of the apoptosis inducer Staurosporine and inhibitor Q-VD-OPh on the density of apoptotic cells and on cell area distributions.**

(A–F′, H-I’) Projections of confocal imaging stacks of fixed epidermis of 20-day starved juvenile polyps labelled for cleaved Caspase substrate (CCS; A-D’, H-I’), phosphorylated ERK1/2 (pERK1/2; E-F’) and F-actin (Phalloidin, A’-I’) after incubation in DMSO (control), Staurosporine (6 hours) or Q-VD-OPh (2 hours). As expected, Staurosporine incubation leads to a strong, concentration-dependent increase in CCS+ apoptotic cells (A-A’) and a broad increase in pERK1/2 signal (E-F’), while Q-VD-OPh treatment strongly reduces CCS+ apoptotic cells across the epidermis (H-I’). (G, J) Violin plots show the distribution of mean cell areas across larger cells in the epidermis after 2 hours of 0.5% or 1% DMSO (control), 0.5μM Staurosporine or 100μM Q-VD-OPh treatments at 5 days post-refeeding (dpr) or 20 dpr. Mean cell areas among larger cells tended to decrease after Staurosporine treatment, which was statistically significant only at 20dpr. The mean cell areas were not significantly affected by Q-VD-OPh. Means represented as black full line and medians as red dashed lines. Single data points represent mean large cell area per animal. Significance codes for model adjusted P-values: * $P<0.05$, ns = non-significant. Abbreviations: dpr: days post-refeeding, Staurosp.: Staurosporine. Scale bars: 20 μm.

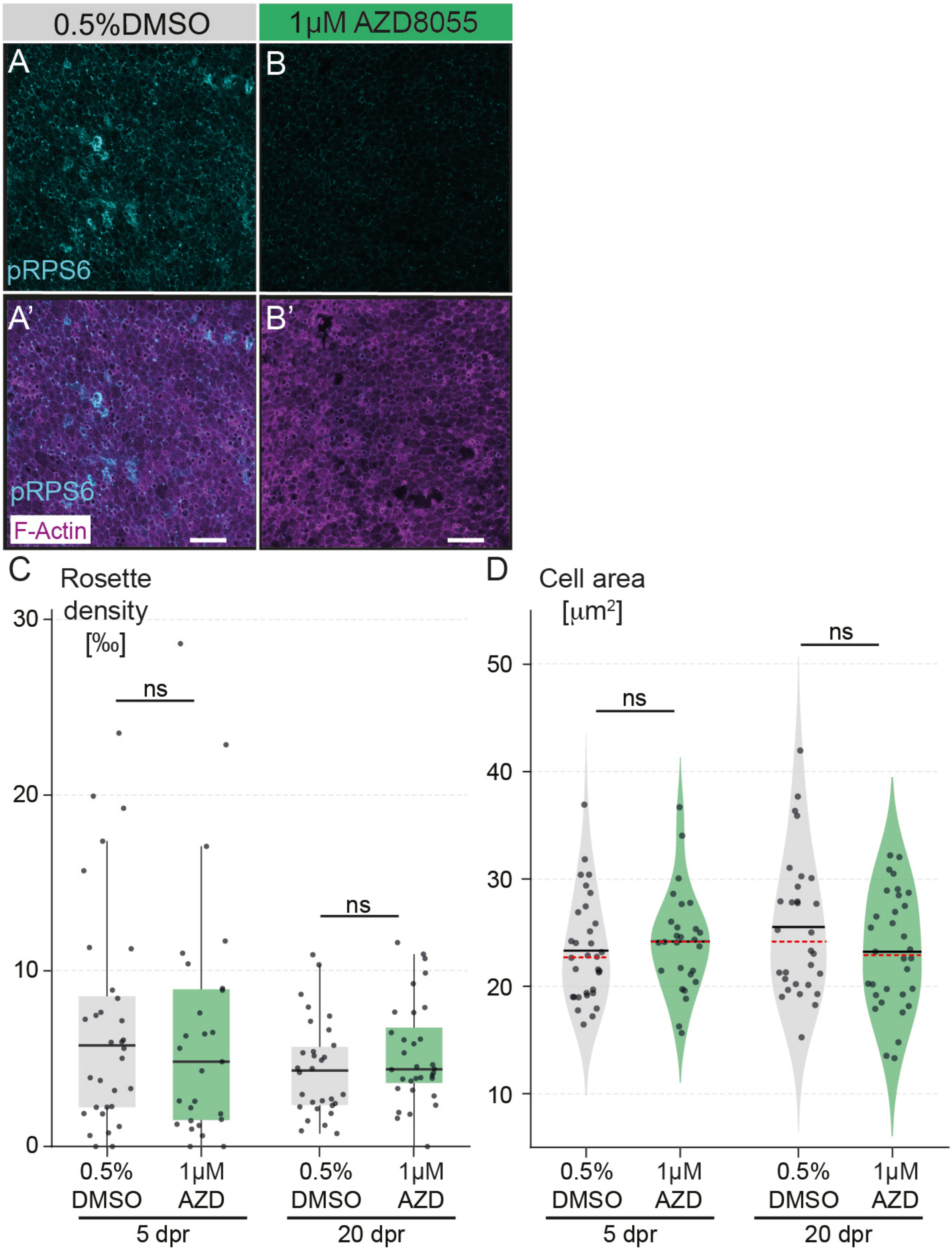


**Fig. S9: Inhibition of TOR signalling does not significantly**

(A-B') Projections of confocal imaging stacks of fixed epidermis of 5-days starved juvenile polyps labelled for phosphorylated ribosomal S6 protein (pRPS6) and F-actin (Phalloidin) after 2 hour-long 0.5% DMSO (A-A') or 1 μM AZD8055 (B-B') treatment. Representative image showing that phosphorylation of RPS6 becomes undetectable in the juvenile body wall epidermis after TOR signalling inhibition by AZD8055. Scale bars: 20 µm. (C-D) Box plots (C) and violin plots (D) show the distribution of rosette densities (C) or cell areas of larger cells (D) in the epidermis after 2 hours of 0.5% DMSO (control) or 1µM AZD8055 treatments at 5dpr and 20dpr. Inhibition of TOR signalling using AZD8055 had no effects on mean rosette density (number of rosettes per 1000 large cells) or mean cell areas of larger cells. Box plots show median values (black full lines), the first to third interquartile ranges (box), whiskers indicate 1.5x IQR. Single data points represent mean rosette density per animal. In violin plots, means are represented by black full lines and medians by red dashed lines. Data points represent means of large cell area per animal. Significance codes for model adjusted P-values: ns = non-significant. Abbreviations: dpr: days post-refeeding, AZD: AZD8055. Scale bars: 20 µm.
